# Satellitome comparison of two oedipodine grasshoppers highlights the contingent nature of satellite DNA evolution

**DOI:** 10.1101/2021.07.01.450629

**Authors:** Juan Pedro M. Camacho, Josefa Cabrero, María Dolores López-León, María Martín-Peciña, Francisco Perfectti, Manuel A. Garrido-Ramos, Francisco J. Ruiz-Ruano

## Abstract

**Background:** The full catalogue of satellite DNA (satDNA) within a same genome constitutes the satellitome. The Library Hypothesis predicts that satDNA in relative species reflects that in their common ancestor, but the evolutionary mechanisms and pathways of satDNA evolution have never been analyzed for full satellitomes. We compare here the satellitomes of two Oedipodine grasshoppers (*Locusta migratoria* and *Oedaleus decorus*) which shared their most recent common ancestor about 22.8 Ma ago.

**Results:** We found that about one-third of their satDNA families (near 60 in every species) showed sequence homology and were grouped into 12 orthologous superfamilies. The turnover rate of consensus sequences was extremely variable among the 20 orthologous family pairs analyzed in both species. The satDNAs shared by both species showed poor association with sequence signatures and motives frequently argued as functional, except for short inverted repeats allowing short dyad symmetries and non-B DNA conformations. Orthologous satDNAs frequently showed different FISH patterns at both intra- and interspecific levels. We defined indices of homogenization and degeneration and quantified the level of incomplete library sorting between species.

**Conclusions:** Our analyses revealed that satDNA degenerates through point mutation and homogenizes through partial turnovers caused by massive tandem duplications (the so-called satDNA amplification). Remarkably, satDNA amplification increases homogenization, at intragenomic level, and diversification between species, thus constituting the basis for concerted evolution. We suggest a model of satDNA evolution by means of recursive cycles of amplification and degeneration, leading to mostly contingent evolutionary pathways where concerted evolution emerges promptly after lineages split.

## Background

Satellite DNA (satDNA) was first described by Kit [1] in mouse and guinea-pig DNA with its repetitive nature demonstrated by Waring and Britten [2]. The first model for satDNA evolution was devised by Smith [3], who demonstrated that DNA sequences that are not maintained by natural selection evolve a tandem repeat structure due to unequal crossing-over. Later, theoretical analyses assumed that satDNA evolution usually depends on mutation, unequal crossing-over, and random drift, with purifying selection controlling for excessive copy number [4,5,6,7,8,9,10,11].

Changes in satDNA amount are mainly due to unequal crossing-over, although other mechanisms have been proposed to explain both amplification and spread of satDNA repeats (for review, see Garrido-Ramos [12]). Walsh [13] proposed the replication of extrachromosomal circles of tandem repeats by the rolling-circle mechanism and reinsertion of replicated arrays as a powerful satDNA amplification process, a mechanism for which Cohen et al. [14,15] have found some support. Additionally, transposition may operate in satDNA emergence and amplification [16,17,18,19]. Ultimately, replication-slippage might be an amplification process [10,13], mainly involved in lengthening satellite monomers from basic shorter ones [20].

To explain the conservation of satellite sequences over long evolutionary periods, Fry and Salser [21] suggested the Library Hypothesis. According to this hypothesis, a group of related species should share a common library of satDNA sequences that mostly show quantitative differences among species due to differential amplification. Therefore, a given member of the library may appear as an abundant satDNA, while others remain at low amounts and technically undetectable. Now we know that the former can be visualized by FISH and the latter discovered by next-generation sequencing [22]. Fry and Salser [21] suggested that an essential step in the evolution of some satDNA families may be the acquisition of a biological function, in which case natural selection would conserve its sequence for long evolutionary periods [23,24,25].

There are some examples of satDNA persisting for long, i.e., more than 40 Ma (see Arnason et al. [26]; Garrido-Ramos et al. [27,28]; de la Herrán et al. [29,30]; Mravinac et al. [31,32]; Robles et al. [33]; Cafasso and Chinali [34]; Chaves et al. [35]). Whereas the conservation of functional satDNA repeats is explained by purifying selection (see references above), the persistence over time of other satDNA arrays lacking apparent function might be simply due to chance events [8,9,13,37]. Therefore, whether satDNA conservation in two or more species is just chance or due to selective events remains unanswered.

Dover [37,38] suggested unequal crossing-over, gene conversion, and transposition as molecular drive mechanisms for the concerted fixation of paralogous variants, which operate independently of natural selection and drift. Recently, this evolutionary pattern has been replaced by the birth-and-death model in the case of coding multigene families [39,40]. Concerted evolution implies that paralogous copies are more homogenized than orthologous ones when two species are compared. SatDNA families comprise thousands or millions of copies of non-coding paralogous repeat units, frequently arranged in many short arrays spread at different genomic locations [17,22,41,42,43,44,45], so that fixation is improbable in these conditions. In fact, although concerted evolution is the predominant pattern for satDNA evolution, non-concerted evolution has also been reported and explained through various factors such as life-history, population, location, organization, number of repeat-copies, or functional constraints (for review, see Garrido-Ramos [12,44]). However, the ultimate causes for concerted or non-concerted patterns are still unknown.

In this paper, we compare the full catalogue of satDNA families (i.e., the satellitome) between two grasshopper species belonging to the subfamily Oedipodinae, *Locusta migratoria* (Lmi) and *Oedaleus decorus* (Ode), which diverged 22.81 Ma [45]. We show the presence of about one-third of orthologous satDNA families whose sequence comparison pointed to mutation and drift as the main drivers of satDNA evolution. We also got estimates of nucleotide turnover rate at the level of consensus sequences (consensus turnover rate, CTR), using 20 orthologous pairs present in both species, and found that they were highly variable and depended on the history of satDNA amplifications. We also analyzed repeat landscapes and developed indices for satDNA homogenization and degeneration and an index for concerted evolution, which may be useful for future research. Also, we propose a general model for satDNA evolution and suggest that the evolution of these sequences constitute a good example of contingent evolution (see Blount et al. [46]).

## Results

### One-third of satDNA families showed sequence homology between species

The range of variation for repeat unit length (RUL) was 8-400 bp for the 60 satDNA families found in *L. migratoria* and 12-469 bp for the 58 families found in *O. decorus*. For subsequent analyses we included only those satDNA families showing more than 100 copies, which excluded the four least abundant satDNAs in *L. migratoria* (Additional file 1: Table S1). After comparing the consensus sequences of all satDNA families present in both species, we found that 21 families in *O. decorus* showed homology with 20 in *L. migratoria* (Additional file 1: Table S2). We assume that these sets of satDNAs showing some sequence identity were already present in the most recent common ancestor of these two species (dated about 22.81 Ma) and thus belonged to the ancestor satDNA library. Therefore, these homologous sets constituted 12 orthologous superfamilies (OSFs) including 31 and 44 subfamilies in *O. decorus* and *L. migratoria*, respectively (Additional file 1: Table S2). On the other hand, the non-shared satDNA families (37 in *O. decorus* and 36 in *L. migratoria*) could have arisen *de novo* after both lineages split, or else they were lost in one of the species.

Between species comparison of basic satellitome features (Table 1) revealed that shared satDNAs did not show significant differences between species for RUL, A+T content, and abundance, but divergence was lower in *L. migratoria*. However, the non-shared satDNAs showed higher RUL and abundance in *O. decorus*. Within species comparisons between shared and non-shared satDNAs failed to show differences in *O. decorus*. In *L. migratoria*, however, the shared satDNA families showed higher RUL, A+T content and abundance, and lower divergence, than the non-shared ones (Table 1). Taken together, these results revealed the presence of many satDNA families showing short monomers among the non-shared ones in *L. migratoria* which also showed lower A+T content and abundance, but higher divergence than those shared with *O. decorus*.

**Table 1.**
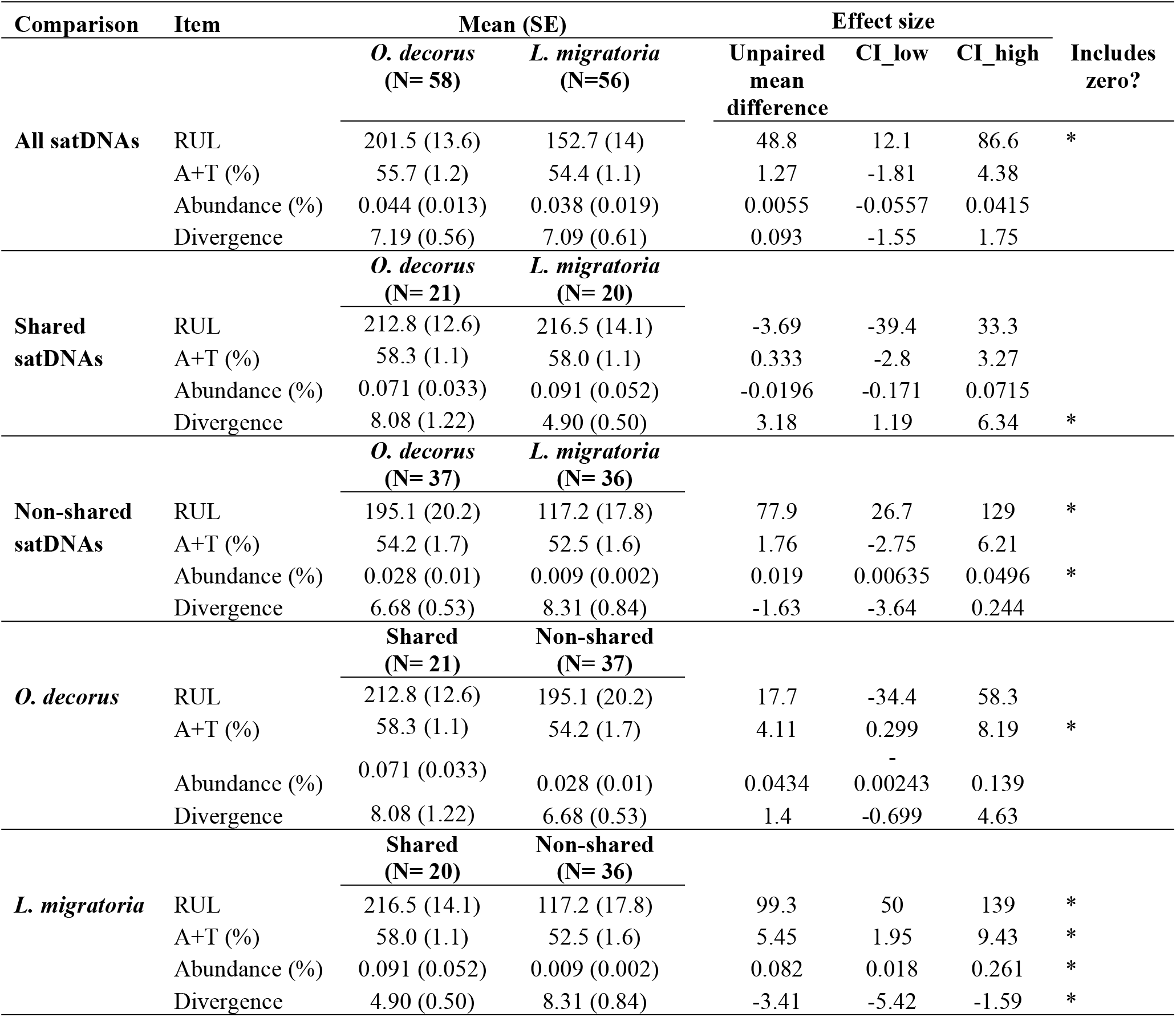
Comparison of satellitome characteristics between *O. decorus* and *L. migratoria* (Southern Lineage), by means of estimation graphics using DABEST (Ho et al. 2019). 95% CI= Confidence interval. RUL= Repeat unit length. * means that 95% CI does not include the zero value.

### Tandem structure and association with other repetitive elements

The quantification of homogeneous and heterogeneous read pairs allowed estimating the degree of tandem structure (TSI) for each satDNA family (Additional file 1: Table S1). The annotation of the heterogeneous read pairs allowed identifying other genomic elements adjacent to satDNA (Additional file 1: Table S3). This revealed that LmiSat03-195 (TSI= 99.7%) was associated with LINEs in 57 out of the 100 heterogeneous read pairs observed. However, only 2% of the 1,356 heterogeneous read pairs showed association with LINEs for its orthologous OdeSat02-204 (TSI= 95.9%), suggesting that association with LINEs occurred only in *L. migratoria*. Likewise, OdeSat17-176 and LmiSat02-176 showed association with Helitron TEs in 93% and 76% of the 2,379 and 1,356 heterogeneous read pairs observed, respectively. Bearing in mind that the sequence of the LmiSat02-176 repeat unit shows homology with Helitron TEs (Ruiz-Ruano et al. 2016), the high frequency of association with Helitron observed for OdeSat17-176 and the low TSI (11.1%) suggest that most units detected for this satDNA were part of the TE itself and are not in tandem (i.e., 1-TSI= 88.9%). However, LmiSat02-176 showed high TSI (94.7%) and lower association with the TE (76%), suggesting that this satDNA arose from this TE, but it also constitutes an independent entity which has reached quite long arrays in *L. migratoria* (longer than 20 kb in the MinION reads). The FISH pattern of both satDNAs (see below) reinforced this conclusion, as OdeSat17-176 yielded no hybridization signals (Table 2), whereas LmiSat02-176 showed pericentromeric bands on six chromosome pairs (see Ruiz-Ruano et al. [22] and Additional file 1: Table S1).

**Table 2.**
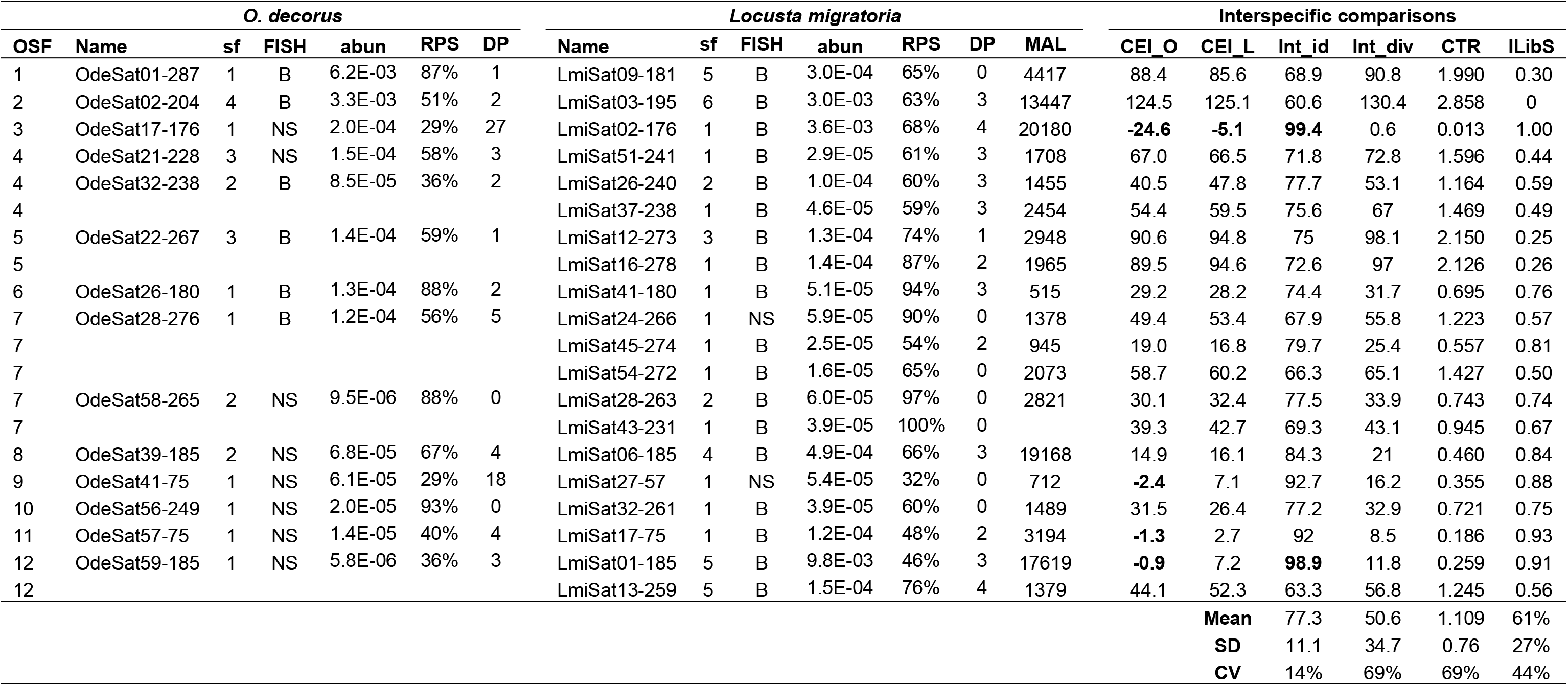
Characteristics of the orthologous satDNA families analyzed in *O. decorus* (14) and *L. migratoria* (20). Each row includes one Ode and one Lmi satDNA families showing homology between them. Note that some Ode families showed homology with two or three Lmi ones. OSF= Orthologous superfamily, sf= number of subfamilies, FISH= FISH pattern (B= banded, NS= no signal), abun= abundance (% of the genome), RPS= Relative peak size, DP= DIVPEAK, MAL= Maximum array length observed in MinIon reads of L. migratoria, CEI= Concerted evolution index (L= *L. migratoria*, O= *O. decorus*), Intid= Interspecific sequence identity (%), Intdiv= Interspecific divergence, CTR= Consensus turnover rate, ILibS= Incomplete library sorting. Negative CEI values and Int_id>95% are remarked in bold type letter. See Table S4 to complete data with repeat unit length, A+T content, divergence (%), peak size, kurtosis of the repeat landscape, tandem structure index and Gibbs free energy of the secondary structure.

### A same orthologous satDNA may show different FISH patterns at intra- and interspecific levels

FISH analysis for 14 OdeSat families, which showed homology with 20 LmiSat ones, revealed that six OdeSats displayed conspicuous bands on chromosomes (B-pattern from hereafter). In contrast, the eight remainders failed to show FISH signal (NS-pattern from hereafter), of which seven showed the B-pattern in *L. migratoria* (Table 2). This revealed that a same OSF may show FISH signals in one species but not in a close relative.

To search for molecular differences between satDNAs showing the B- and NS-patterns, we analyzed MinION long reads in *L. migratoria* to score the maximum array length (MAL) for each LmisatDNA (Table 2). Even though coverage was very low (0.02x), we found that none of the seven NS families analyzed showed arrays higher than 2,500 bp, whereas almost half of those showing the B pattern did (Gardner-Altman unpaired mean difference= 2930, 95.0%CI: 1540, 4790), and the three orders of magnitude of the difference indicated that satDNAs with the B-pattern have been submitted to more (and extensive) amplification events than those showing the NS-pattern. This difference justifies using the presence of FISH signals as an indication of the degree of satDNA amplification. The fact that 18 out of 20 orthologous satDNA families in *L. migratoria* showed the B-pattern, whereas only six out of the 14 orthologous families analyzed in *O. decorus* showed it, represent the first indication for a higher incidence of satDNA amplifications in *L. migratoria* (RxC contingency test, with 50,000 replicates: P= 0.00562, SE= 0.00077). This result was reinforced by the fact that the 14 OdeSat families included 24 subfamilies whereas the 20 LmiSat ones included 44 subfamilies (Table 2) (Wilcoxon matched-pairs test: z= 2.11, N=12, P= 0.035). As subfamilies represent different amplification events, the former results demonstrate that a same orthologous satDNA may show different amplification trajectories during their independent evolution in different species.

Careful examination of orthologous satDNAs revealed a unique case of no satDNA amplification in both species during the 22.8 Ma of separate evolution, as the LmiSat27-57 and OdeSat41-75 OSF showed the same NS-pattern. Consistently with their low degree of amplification, these two satDNAs showed very low values for tandem structure (TSI: 9% in *O. decorus* and 32% in *L. migratoria*) and homogenization (RPS: 29% and 32%) indices (see next section), indicating poor tandem structure and homogenization (see Table 2 and Additional file 1: Table S4). The remaining OSFs, however, showed amplification in at least one species. One of the most dramatic differences was found for the orthologues OdeSat59-185 and LmiSat01-185, which were the scarcest and the most abundant satDNAs in *O. decorus* and *L. migratoria*, respectively, with the latter showing pericentromeric FISH bands on all chromosomes [22] and OdeSat59-185 showing the NS-pattern (Fig. 2 and Table 2). In fact, seven orthologous satDNA families with the NS-pattern in *O. decorus* showed the B-pattern in *L. migratoria* (Table 2 and Fig. 3).

**Figure 1.**
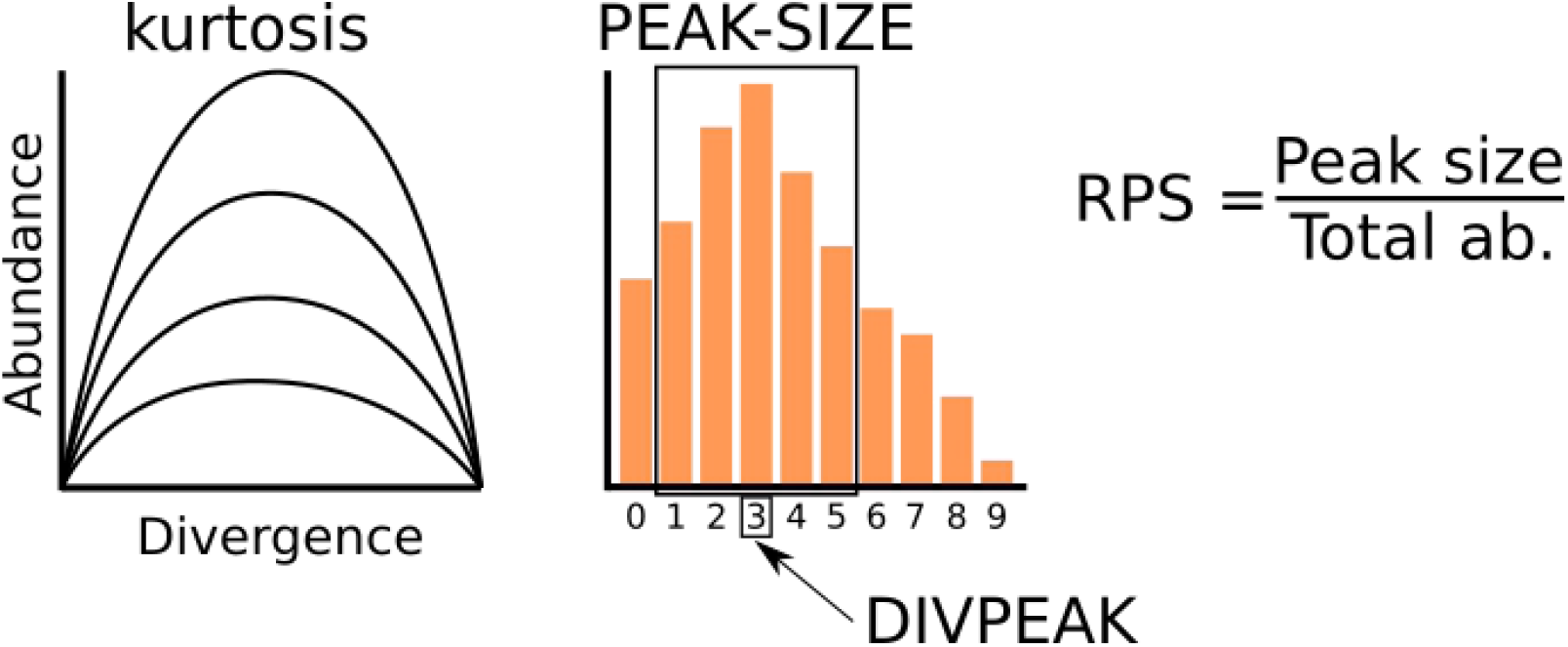
Definition of satDNA parameters in respect to abundance and divergence. The distribution of the abundances of groups of sequences differing by 1% divergence constitutes a repeat landscape (RL). It may be seen as a curve (left) or an histogram (right). In addition of variation in kurtosis, represented by several curves on the left, three properties of satDNA can be defined on RLs: DIVPEAK is the divergence class showing the highest abundance (3% in the histogram); PEAK-SIZE is the sum of the abundances of the five classes included around DIVPEAK, thus constituting the sum of all sequences differing by less than 5%, thus coinciding with our definition of satDNA subfamily; RPS is the relative peak size and represents the fraction of abundance which is included in the 5% amplification peak.

**Figure 2.**
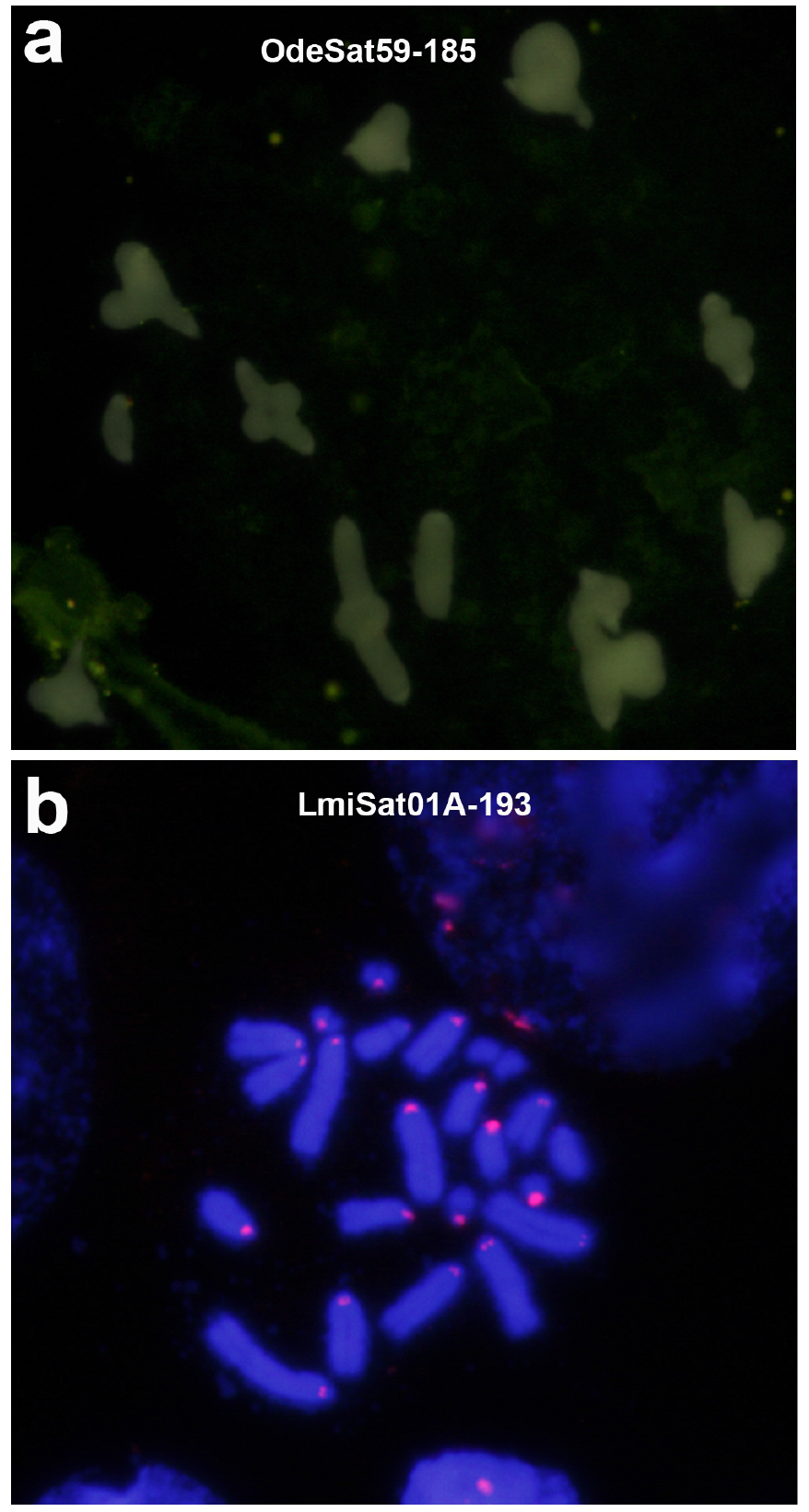
FISH analysis of a pair of orthologous families, belonging to OSF12, in *O.decorus* (a) and *L. migratoria* (b). a) OdeSat59-185 showed no FISH bands on this meiotic metaphase I cell, thus showing the NS pattern. b) LmiSat01A-193 showed conspicuous pericentromeric FISH bands on most chromosomes of this embryo mitotic metaphase cell, thus showing the B-pattern.

**Figure 3.**
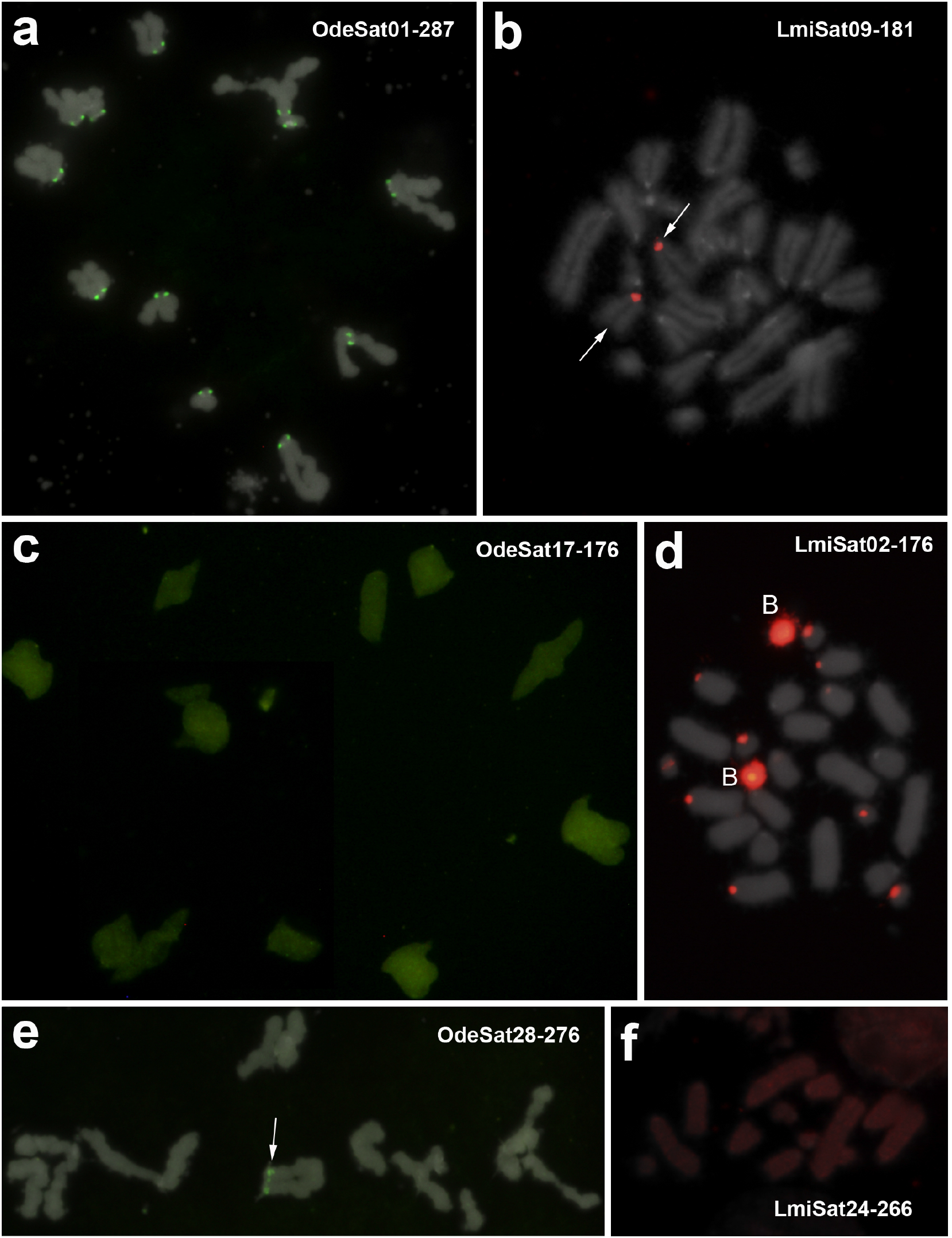
FISH analysis of three pairs of orthologous families in *O.decorus* and *L. migratoria*: showing the B-pattern in both species for OSF1 (a and b), the NS and B patterns, respectively, for OSF3 (c and d), and and the B and NS patterns, respectively, for OSF7 (e and f) (see also Table 2 for satDNA classification into OSFs). a) Presence of pericentromeric FISH bands for OdeSat01-287 on all chromosomes of this meiotic metaphase II cell of *O. decorus*. b) Note the presence of its orthologous family (LmiSat09-181) on a single chromosome pair of this embryo mitotic metaphase cell of *L. migratoria*. c) Absence of FISH bands for OdeSat17-176 in a meiotic metaphase I cell of *O. decorus*. d) Presence of its orthologous LmiSat02-176 on pericentromeric regions of several chromosome pairs and on whole B chromosome length (B) of this embryo mitotic metaphase cell of *L. migratoria*. e) Presence of a pericentromeric FISH band on a single chromosome of the haploid set shown in this meiotic metaphase II cell of *O. decorus*. f) Absence of FISH bands for LmiSat24-266 on the haploid chromosome set shown in this embryo mitotic metaphase cell of *L. migratoria*.

An interesting case was OSF7, where one of the five *L. migratoria* families showed the NS-pattern (LmiSat24-266) whereas the four remaining (LmiSat28-263, LmiSat43-231, LmiSat45-274 and LmiSat54-272) showed the B-pattern (Table 2). Likewise, one of the two *O. decorus* families (OdeSat28-276) showed the B-pattern whereas the other (OdeSat58-265) showed the NS one. This shows that homologous satDNAs can display the NS or B patterns at intra- and interspecific levels. Finally, even those satDNAs with FISH bands in both species showed remarkable differences regarding chromosome location (proximal, interstitial, or distal; see Additional file 1: Table S1). Taken together, these results show that orthologous satDNAs can display disparate chromosome distribution in separate species due to their independent evolution, a fact previously reported in the literature [47,48,49,50]. These differences can range from short arrays being undetectable by FISH, which may eventually serve as seeds for species-specific amplification (as suggested by Ruiz-Ruano et al. [22]), up to long arrays yielding conspicuous FISH bands.

### SatDNA homogenization and degeneration

SatDNA homogenization and degeneration are considered important drivers of satDNA evolution, but their relative importance has been debated. It would thus be desirable to find satDNA parameters being good indices for these two alternative states. To search for a homogenization index, we hypothesized that it should show a high negative correlation with intraspecific divergence. Spearman rank correlation analysis showed that, in both species, RPS (relative peak size, see methods and Fig. 1) showed a very high negative correlation with divergence (measured as K2P) (*r_S_*= -0.9 in both species) (Table 3), which revealed RPS as a good homogenization index. On the contrary, a degeneration index should be negatively correlated with homogenization, and Spearman rank correlations revealed that DIVPEAK (i.e. the divergence value showing the maximum abundance in a repeat landscape, see Fig. 1) showed the highest negative correlation index with RPS in both species (Table 3). This means that the relative size of amplification peaks decreases as satDNA sequences accumulate divergence through mutational decay since the last satDNA amplification (see repeat landscapes in Fig. 2, Additional file 2: Fig. S1 and Additional file 3: Dataset 1).

To ascertain whether satDNA degeneration, measured by DIVPEAK, is associated with any of the satDNA parameters analyzed (RUL, A+T, no. subfam and TSI), we performed Spearman rank correlation analyses, which revealed that RUL was the only satDNA property showing significant correlation with DIVPEAK (Table 3) and it was negative and of similar magnitude as that between DIVPEAK and RPS. This suggests that RUL is an important determinant of satDNA degeneration, with shorter satDNAs degenerating faster. A possible explanation is that short monomers degenerate faster through mutational decay because every point mutation implies a higher proportion of degeneration for short than for long monomers, as if the Muller’s ratchet would have fewer teeth for short than long repeat units and the same number of new mutations would imply a higher number of ratchet’s turns for short repeating units than for long ones.

The analysis of the statistical properties of RPS and DIVPEAK indicated that, in both species, RPS fitted a normal distribution (ODE: χ^2^= 4.45, df= 3, P= 0.215; LMI: χ^2^= 4.78, df= 3, P= 0.189 whereas DIVPEAK fitted an exponential distribution (ODE: χ^2^= 4.55, df= 2, P= 0.103; LMI: χ^2^=4.93, df= 3, P= 0.177). Their scales ranged between 0 and 1 for RPS and between 0 and 27% (within the 0-40% scale of divergence measured here) for DIVPEAK.

To apply these indices to satDNA evolution, we consider that satDNA families follow evolutionary pathways that include recursive cycles of homogenization (through amplification by tandem duplication) and degeneration (through random mutation). After an amplification event, homogenization (measured by RPS) will increase, and degeneration (measured by DIVPEAK) will decrease. As time goes by, with no other amplification events, RPS will decrease and DIVPEAK will move towards higher values. An expected outcome of mutation accumulation is reducing the kurtosis of the repeat landscape (RL) distribution (i.e., curve flattening, Fig. 1 for examples). In fact, kurtosis was correlated negatively with DIVPEAK (Ode: N=58, *r_S_*= -0.80, t= 9.89, P<0.000001; Lmi: N=56, *r_S_*= -0.76, t= 8.58, P<0.000001) and positively with RPS (Ode: N=58, *r_S_*= 0.80, t= 9.68, P<0.000001; Lmi: N=56, *r_S_*= 0.83, t= 10.98, P<0.000001). Kurtosis is thus proportional to RPS, so that highly homogenized satDNAs show leptokurtic RLs whereas highly degenerated ones show platikurtic RLs. Therefore, kurtosis and RPS are expected to be high for recently amplified satDNAs and low for satDNAs that have not been amplified for a long time (see some examples in Fig. 2 and Additional file 2: Fig. S1). Although these parameters do not constitute absolute measures of time, however, they can be useful as measures of “time since the last satDNA amplification”. As satDNA can undergo successive amplifications across evolutionary time, we can also consider RPS and kurtosis as homogenization indices indicating how far is a satDNA from degeneration.

To analyze whether conservation of the orthologous satDNA families in both species was associated with homogenization and degeneration indices, we compared them between the shared and non-shared satDNA families found in each species. In *O. decorus*, the effect size (unpaired mean difference) found between non-shared and shared satDNAs by means of Gardner-Altman estimation plots, revealed no mean differences for RPS (unpaired mean difference= -0.0682, 95.0%CI: -0.159, 0.0348), kurtosis (unpaired mean difference= 0.678, 95.0%CI: -1.62, 5.78) and DIVPEAK (unpaired mean difference= 1.13, 95.0%CI: -0.954, 5.61), indicating similar levels of homogenization and degeneration in both groups. In *L. migratoria*, however, the three indices showed differences between shared and non-shared satDNA families, indicating higher homogenization and lower degeneration for the shared ones (Fig. 4).

**Figure 4.**
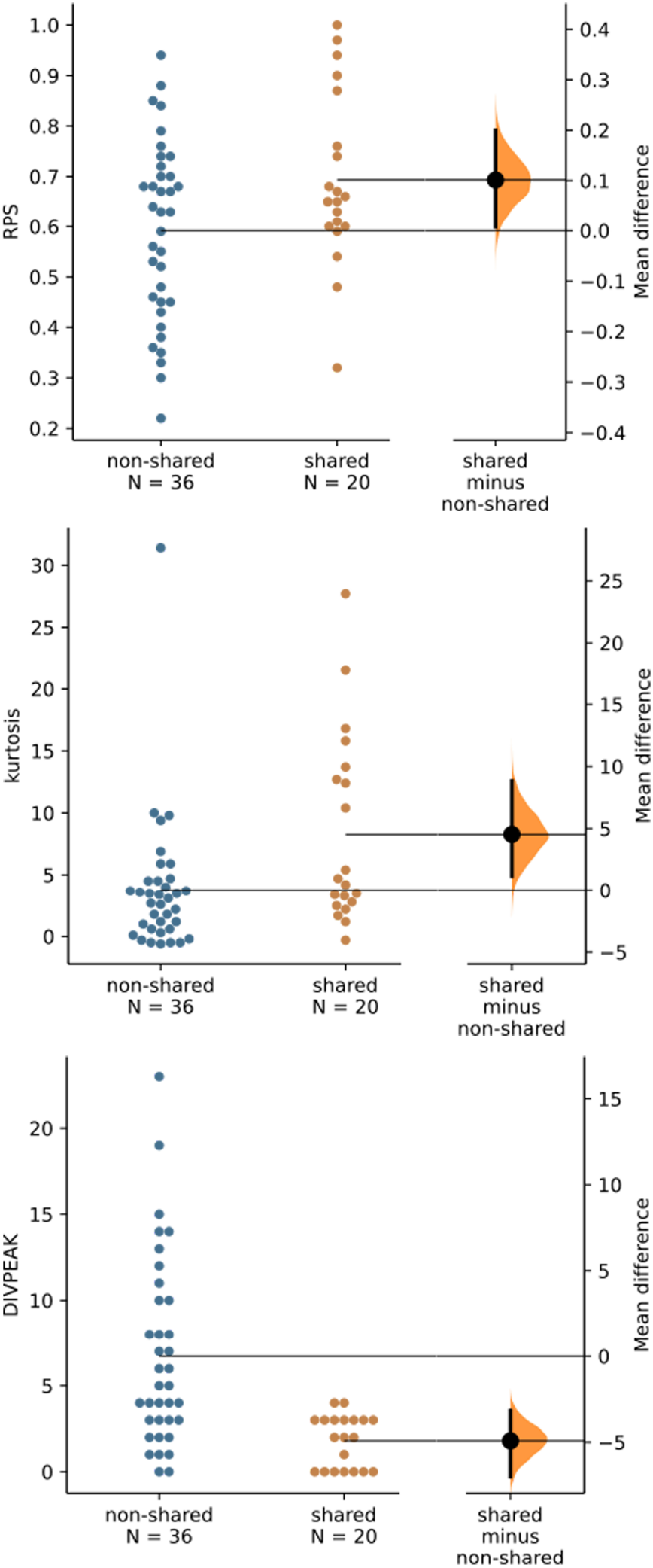
Repeat landscape (RL) and minimum spanning tree (MST) of two orthologous superfamilies of satellite DNA in *O. decorus* and *L. migratoria* (OSF02 and OSF12). a) OSF02 showed the highest consensus turnover rate (CTR= 2.86) found among the 20 values estimated between orthologous pairs of families in both species. Note that OSF02 showed large amplification peaks in both species (green curve in *O. decorus* and red curve in *L. migratoria*) and that the MST showed complete separation of OdeSat02 and LmiSat03 sequences. b) OSF12 showed the lowest CTR estimate (0.26 between OdeSat59 and LmiSat01) and the MST (on the right) reveals that the consensus DNA sequences of these two satDNA families showed only two differences. Also note in the RL (on the left) that the OdeSat59 curve is very close to zero, as this is the satDNA family in *O. decorus* showing the lowest abundance, indicating that OSF12 is represented in this species as relict remains which, by chance, almost coincide in consensus sequence with the most abundant subfamily in *L. migratoria* (LmiSat01A), thus evidencing extreme incomplete lineage sorting (see other cases in Additional file 2: Fig. S1).

### Amplification explains the concerted evolution of satDNA

*O. decorus* and *L. migratoria* shared their most recent common ancestor 22.81 Ma, on which basis we could perform estimations of interspecific rates of turnover in the consensus sequences (CTR). For this purpose, we compared the consensus DNA sequences of 20 pairs of orthologous satDNA, representing half of the 40 estimations that could be done at family level (see Additional file 1: Table S2). The values obtained for CTR in the 20 orthologous pairs ranged from 0.013% (between LmiSat02-176 and OdeSat17-176) to 2.86% (between LmiSat03-195 and OdeSat02-204) nucleotidic changes in their consensus sequences per million year (mean= 1.11%, see Table 2), with two orders of magnitude between the extreme values.

To search for possible causes for such an extreme variation in the observed rates, we performed forward stepwise multiple regression of CTR (dependent) on four factors related to satDNA amplification: for each species, the number of subfamilies per satDNA family (subfam), the absolute number of copies included in the 5% divergence peak (peak-copies), RPS, and TSI. The results revealed that only three out of the eight factors entered a model that explained 85% of the total variance in CTR, with Ode_subfam explaining 56.4%, Ode_peak_copies explaining 25.7%, and TSI_Ode explaining only a nonsignificant 2.8% (Table 4). Variance inflation factors of this regression analysis ranged between 1.07 and 3.01 indicating the absence of multicollinearity. Likewise, the standardized residuals of this regression fitted a normal distribution (Shapiro-Wilks test: W= 0.97, P= 0.82). Finally, partial correlations were 0.85 for Ode_subfam, 0.76 for Ode_peak_copies, and 0.40 for TSI_Ode, whereas they were much lower for the five factors failing to enter in the model (from -0.25 to -0.02).

As we defined satDNA subfamilies by sharing 95% or higher sequence identity, i.e., up to 5% divergence, which was exactly the same figure used to define RPS and DIVPEAK on RLs, we consider that the number of subfamilies actually represents the number of independent amplification events being apparent within each family, as it also coincides with the number of different consensus sequences per family. As peak-copies represents the total number of repeat units in the amplification peak, we can infer that the rate of nucleotide change estimated from consensus sequences (CTR), which is positively correlated with the two former parameters, roughly represents the rate of nucleotide changes driven by satDNA amplification to be part of the consensus sequence. It was remarkable that only *O. decorus* variables entered in the stepwise multiple regression model, as it is the species showing the lowest number of subfamilies (31 versus 44 in the 12 OSFs, as a whole, and 24 and 44 in the 14 orthologous pairs analyzed) and thus showed fewer amplification events, suggesting that CTR value is limited by the species showing fewer amplification events. We thus conclude that the same molecular mechanism, i.e., satDNA amplification, causes intraspecific homogenization and interspecific diversification, thus explaining the concerted evolution pattern of satDNA.

### Most satDNA families showed concerted evolution in both species

Concerted evolution predicts that CEI>0, and this was met for 16 orthologous pairs, the four exceptions being the OdeSat17-LmiSat02 pair and three satDNA families in *O. decorus* (OdeSat41, OdeSat57, and OdeSat59) where CEI<0 thus showing signs of non-concerted evolution (Table 2). Remarkably, these four OdeSats failed to display FISH bands, suggesting that poor amplification might be related with non-concerted evolution. In both species, CEI was positively correlated with RUL (Ode: *r_S_*= 0.70, N= 14, t= 3.4, P= 0.0051; Lmi: *r_S_*= 0.56, N= 20, t= 2.83, P= 0.011) and RPS (Ode: *r_S_*= 0.73, N= 14, t= 3.67, P= 0.0032; Lmi: *r_S_*= 0.68, N= 20, t= 3.88, P= 0.0011) but not with A+T content (P>0.05 in both species). In addition, CEI was positively correlated with TSI in *O. decorus* (*r_S_*= 0.78, N= 14, t= 4.26, P= 0.0011) but not in *L. migratoria* (*r_S_*= 0.43, N= 20, t= 2.04, P= 0.056). Finally, in *O. decorus*, CEI was higher in the six satDNAs showing the FISH B-pattern than in the eight showing the NS-pattern (unpaired mean difference= 2.63; 95% CI: 0.883, 5.36).

These results indicate that satDNAs displaying longer monomers, higher levels of homogenization and the FISH B-pattern show higher indices of concerted evolution. Exceptional non-concerted patterns were observed for satDNA families showing a low number of amplifications since all showed a single subfamily in both species.

### The persistency of satDNA in these two species was not associated with functional constraints

Several sequence features have hitherto been associated with a variety of putative satDNA biological roles, the most relevant being centromere function. We searched for short internal repeats within each satDNA family’s consensus sequences since these repeats have been associated with sequence function. We found no direct repeats within the sequence span of any satDNA sequence. On the contrary, it was common to find short inverted repeats in all satDNA families that might facilitate non-B DNA conformations such as stem-loops and cruciform structures, but they were found in both shared and non-shared satDNA families.

To ascertain whether Gibbs free energy (dG) of satDNA sequence depends on some satDNA properties, we performed forward stepwise regression, in each species, with dG as dependent variable and RUL, A+T, sharing status and degeneration status (DIVPEAK) as independent factors. In Ode, the regression model explained 67% of the variance in dG (59% by RUL, 5% by A+T, and 3% by DIVPEAK). The correlation was negative with RUL and positive with the two other factors. In *L. migratoria*, the result was highly similar, except that DIVPEAK did not enter in the model, but the dG variance explained was higher, reaching 83% (79% by RUL and 4% by A+T). As higher free energy values correspond to lower dG values, the former results indicate that free energy of satDNA sequence depends positively on RUL, as it determines the likelihood of autopairing, and, at lower extent, also depends on two other sequence properties influencing the number of hydrogen bonds in the double helix, as higher A+T content implies more A-T pairs and fewer hydrogen bonds, thus lower free energy, whereas higher DIVPEAK indicates higher mutational decay that might difficult autopairing thus decreasing the number of hydrogen bonds. The fact that DIVPEAK of the shared satDNAs was higher in *O. decorus* than *L. migratoria* (paired mean difference= 2.6, 95.0%CI: 0.55, 6.8) is consistent with their higher degeneration in *O. decorus*.

We found that most of the shared satDNA families failed to show a propensity to acquire stable curvatures (Additional file 1: Table S1), even though the curvature-propensity plots contained a peculiar maximum in some of them. However, the magnitude of these peaks (11 to 13 degrees/10.5 bp helical turn) was far from the values calculated for other highly curved motifs [51,52]. Most intriguingly, these peaks were similar for satDNAs showing the NS or B FISH patterns or, in the latter case, whether they were located on pericentromeric regions or not. In total, only 11 (7 in *L. migratoria* and 4 in *O. decorus*) out of the 34 shared satDNA families showed curvature propensity, all showing RUL≥185 bp. They belonged to five different OSFs, three of which showed curvature propensity in both species, whereas the two remaining showed it in only one species, suggesting that this property does not depend only on RUL, which was highly similar in both species for these satDNA families.

We also analyzed curvature propensity for the non-shared satDNAs, and none of them showed it to a large degree. Notwithstanding, as observed for shared satDNAs, a few families (one in *L. migratoria* and five in *O. decorus*) showed a conspicuous peak of magnitudes between 11 to 14 degrees/10.5 bp helical turn. It has been suggested that DNA curvature may be involved in the recognition of DNA-binding protein components of the heterochromatin [53]. Our results show that curvature propensity is not differentially frequent or relevant in the 34 shared satDNAs analyzed in both species, compared with the non-shared ones. Therefore, we believe that curvature propensity is not a relevant feature of satDNA or the cause for satDNA conservation in these two species.

Finally, we searched for the presence of short sequence motifs common to the shared satDNA families in both species. We isolated individual monomers from each satDNA family and calculated nucleotide diversity (π) per position (not shown). We did not find conserved motifs in these satDNAs, irrespectively of their FISH pattern or chromosomal location.

Taken together, these results show that, in these two species, there is no sequence conservation for pericentromeric satDNAs, which also lack significant sequence signatures other than A+T richness and repeat length. On the other hand, all putative functional signatures analyzed here were not more frequent in the shared satDNAs than in the non-shared ones. We interpret this as evidence that satDNA conservation is mostly a contingent event. This conclusion is logically conditioned by data and methodology limitations, such as testing based just on sequence data and genomic location, and using a long time scale.

### Incomplete sorting of the satDNA library

The satellitomes of relative species show sequence homology for a fraction of their satDNA families, which is the best support for the satDNA library hypothesis [21]. Joint analysis of RLs and MSTs revealed interesting properties of the satDNA library (Fig. 2 and Additional file 2: Fig. S1): i) OdeSat02A and LmiSat03A were the two OSF02 subfamilies showing the highest amplification peaks in the RLs (Fig. 5a, plot on the left), and they also showed the highest CTR observed among all those analyzed here (2.86% per Ma). Remarkably, the MST plot for all subfamilies and families comprising OSF02 revealed complete sorting per species for this component of the library (Fig. 5a, right). ii) On the other hand, OSF12 included two families in *L. migratoria* (LmiSat01 and LmiSat13) which were fully sorted in the MST (Fig. 5b, right), whereas the single *O. decorus* family (OdeSat59) was remarkably similar to LmiSat01A, with only two nucleotidic differences in their sequence, which is lower than those shown by the four other *L. migratoria* subfamilies with LmiSat01A. This illustrates an extreme case of incomplete library sorting (ILibS) and the second lowest CTR value (0.26% per Ma). Other OSFs showed intermediate situations. For instance, OSF04 showed CTR values between 1.16 and 1.60 and their MST revealed the existence of ILibS, with OdeSat32A being connected with three different LmiSats (37A, 26A and 51A), the latter being placed betwee OdeSat32A and OdeSat21A (see Additional file 2: Fig. S1a). On the contrary, OSF5 (Additional file 2: Fig. S1b) showed high CTR values (>2% per Ma) and complete library sorting, with the satDNAs properly separated between species. Finally, OSF07 showed CTRs between 0.56 and 1.43 and apparent ILibS, with high level of intermixing between the satDNAs of both species (Additional file 2: Fig. S1c). Taken together, these observations suggest that CTR values are inversely associated with the level of ILibS. On this basis, we used the maximum CTR value (maxCTR= 2.86) as reference to estimate the degree of ILibS as one minus the quotient between CTR_i_ and maxCTR (see Table 2). This indicated that the satDNA library of *O. decorus* and *L. migratoria* shows, on average, 61% of incomplete sorting after 23 Ma. Finally, the fact that the four OdeSats showing the non-concerted pattern were those showing the highest ILibS figures (0.88-1), whereas ILibS values up to 0.84 corresponded with patterns of concerted evolution (see OSF8 in Table 2), suggested the possible existence of a threshold for ILibS (between 0.84 and 0.88) below which satDNA evolution is concerted.

**Figure 5.**
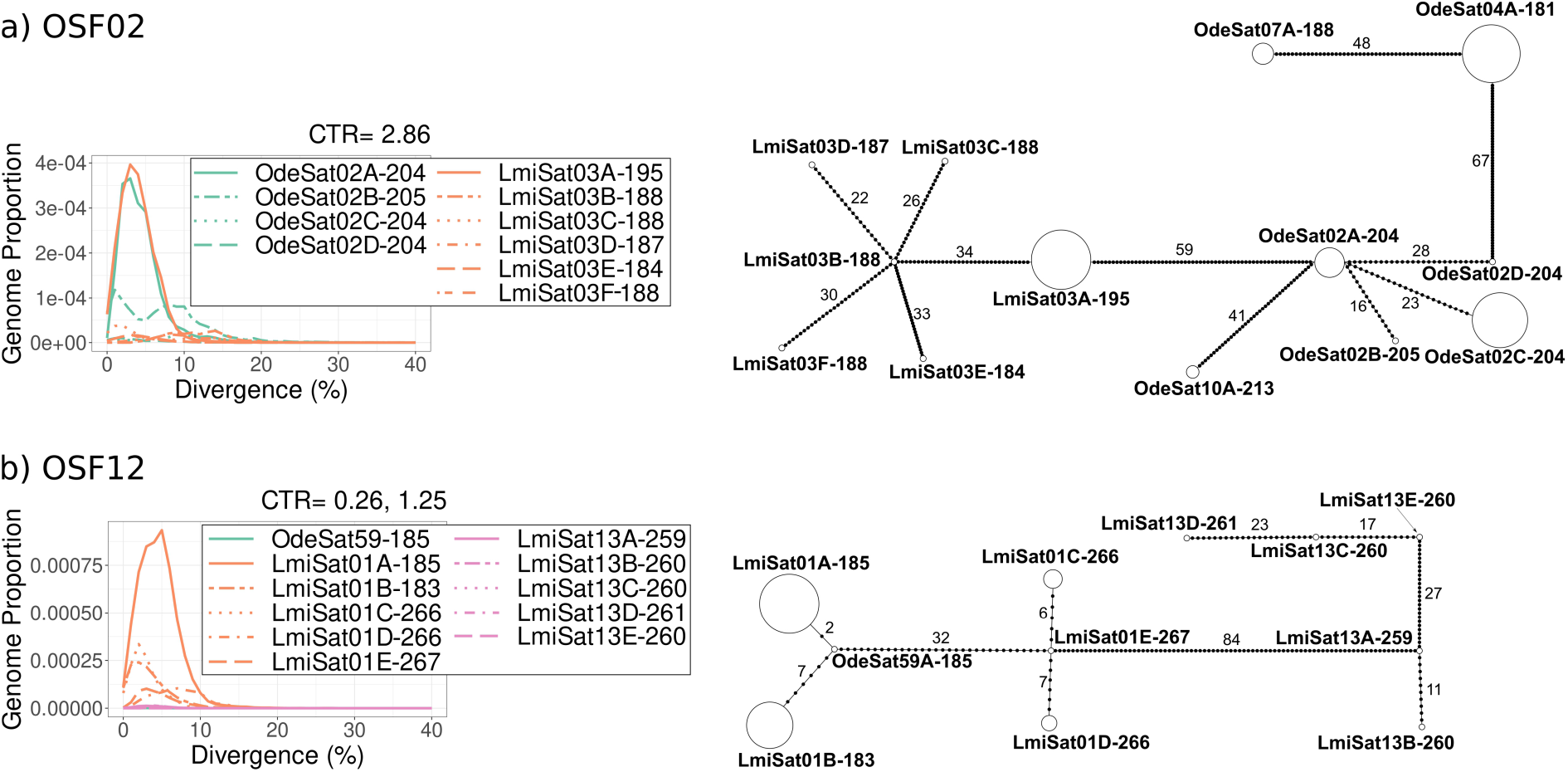
Gardner-Altman plots comparing RPS, kurtosis and DIVPEAK between the *L migratoria* satDNA families being shared or non-shared with *O. decorus*. Note that shared satDNAs showed higher homogenization (higher RPS and kurtosis) and lower degeneration (5% effect size for mean difference in DIVPEAK) than non-shared ones, suggesting most recent amplification of the shared ones.

## Discussion

### SatDNA evolution is mostly contingent

Comparative analysis of the satellitome in the grasshoppers *O. decorus* and *L. migratoria*, two species belonging to the Oedipodinae subfamily, which shared their most recent common ancestor about 23 Ma, gave us a chance to take a look into satDNA library evolution during this period. We assume that the 41 satDNA families (20 in *L. migratoria* and 21 in *O. decorus*) that showed sequence homology between species belong to 12 orthologue groups already present in the ancestor library, which have been conserved up today. However, the remaining 84 families (36 in *L. migratoria* and 37 in *O. decorus*) could represent either remnant satDNAs conserved in only one species or satDNAs arisen *de novo* during the separate evolution of these species. To distinguish between these two possibilities, it is necessary to analyze other oedipodine species. The occurrence of a species-specific profile of satDNAs resulting from differential amplifications and/or contractions from a pool of sequences shared by related genomes is a prediction of the library hypothesis of satDNA evolution with the subsequent replacement of one satDNA family for another in different species [21]. By analogy with incomplete lineage sorting (ILS) in phylogenetic studies, satDNA amplifications and/or contractions between close relative species may yield a pattern of incomplete library sorting (ILibS). We have detected here this phenomenon using consensus sequences, but the use of physical sequences would yield even higher rates of ILibS.

The library hypothesis predicts the residual retention of low-copy counterparts of the dominant satDNA of one species in the other [21]. For instance, OdeSat02A-204 and LmiSat03A-195 have been independently amplified in both species, reaching among the highest genomic abundances in both species, and showed the highest CTR and extensive diversification, with four subfamilies in *O. decorus* and six in *L. migratoria* (see Fig. 5a). In addition, a joint MST for OSF02 (to which both satDNA families belong) revealed the absence of ILibS as all satDNA families and subfamilies appeared well separated between species in the MST (see Fig. 5a). Conversely, the consensus sequences of LmiSat01A-185 and OdeSat59-185 only differed in two positions, thus showing higher interspecific similarity than that found, at intraspecifical level, between the five *L. migratoria* subfamilies (see Fig. 5b), thus constituting an extreme example of ILibS. The high similarity in the consensus sequences of OdeSat59A and LmiSat01A cannot be explained by functional conservation because only the latter shows FISH bands on centromeric regions of all chromosomes thus probably playing a centromeric function in *L. migratoria*, whereas OdeSat59A is the most scarce satDNA found in *O. decorus* thus being only a relic. Likewise, while OdeSat01-287 is the most abundant satDNA in *O. decorus*, its orthologous (LmiSat09-181) is a relict in *L. migratoria*. We thus believe that the observed sequence similarity between OdeSat59A and LmiSat01A might be due to chance convergence, as the likelihood of nucleotide coincidence in each position of the consensus sequence is a function of the relative frequency of the four possible nucleotides in each species, thus being a probabilistic issue.

Our estimates of ILibS from CTR values indicated that the satDNA libraries of *O. decorus* and *L. migratoria* still show 61% of incomplete sorting after 23 Ma of independent evolution, i.e. about 39% of complete sorting (1.7% per Ma). This extreme cohesiveness of the satDNA library is due to the highly paralogous nature of these genomic elements, with thousand copies evolving at once, independently in both species, through point mutation, amplification (tandem duplication) and drift (see below). This 39% expresses only part of library divergence, as the maximum divergence would be reached when all homology signals between satDNAs in both species would have been erased, as in the case of the non-shared ones, whereas the satDNAs belonging to OSF02 are still recognized as homologous between species even with 100% library sorting. Anyway, the ILibS parameter of a given OSF (or orthologous pair of satDNAs) inversely indicates its possible utility for phylogenetic analysis.

Another prediction of the library hypothesis is that the appearance of satDNA families would usually represent amplification of one of the satellites already present at a low level in the library, rather than actual *de novo* appearance. It is not easy to know if any of the non-shared satDNA families actually arose *de novo.* However, in *L. migratoria*, the lower RUL of non-shared satDNAs suggests that the satellitome of this species might harbor some *de novo* arisen short satellites, in consistency with an evolutionary trend towards increasing monomer length and complexity, suggested by theoretical [54] and experimental [20,27,29,55] work.

Our estimates of CTR by the comparison of 20 orthologous pairs of satDNA families indicated that it was 1.11% per Ma, which implies that two satellites can diverge by more than 50% in about 50 Ma. This explains why *L. migratoria* and *O. decorus*, belonging to the Acrididae family do not share a single satDNA family with *Eumigus monticola* [56], a grasshopper belonging to the Pamphagidae family, as these two orthopteran families shared their most recent common ancestor about 100 Ma [45]. Along with the stochastic nature of satDNA loss or gain during evolution, sequence changes at the mentioned rate will make unrecognizable a satDNA family after 100 Ma of separate evolution within the genomes of different species, which contrasts with the case of some other satDNAs preserved for more than 60 Ma [28,30,31,34] or even more than 100 Ma [29,33].

Our results suggest that the same OSF may be involved in the centromeric function in a given species but not in a close relative species. According to Melters et al. [57], the most abundant satDNAs in a genome are most likely involved in the centromeric function. Another feature suggesting this fact is satDNA location on pericentromeric regions of all chromosomes. Therefore, LmiSat01-185, OdeSat01-287 and/or OdeSat02-204 are the best candidate families in these species since all meet the two conditions. However, all three satDNAs showed orthologous families in the other species displaying much more limited chromosome distribution, suggesting that one or both species have replaced the centromeric satDNA during the last 22.8 Ma. No significant track of signatures such as conserved motifs or sequence mediated specific stereo-spatial features were found for these or any other pericentromeric satDNAs found in these species. We thus believe that, in the absence of other evidence, contingent facts such as the opportunity to be in the right place when amplified might be responsible for centromeric satDNA turnover. Zhang et al. [58] also revealed rapid divergence for centromeric sequences among closely related *Solanum* species and suggested that centromeric satellite repeats underwent boom-bust cycles before a favorable repeat became predominant in a species. Indeed, there are species such as chicken [60], common bean [60], or pea [61] that contain different satDNAs in different centromeres.

Whether a given satDNA is conserved for long due to functional reasons is an open question. Fry and Salser [21] suggested that an essential step in the evolution of a specific satDNA family may be acquiring a biological function. However, persistence over time of a satDNA might also be explained in terms that do not depend on natural selection [8,9,10,13,36]. Our results were consistent with this latter view. No conserved functional motifs were found within the monomers of every grasshopper satDNA analyzed as has been found in other satDNAs such as human centromeric satDNA [62,63,64,65]. On the other hand, short dyad symmetries within satDNA repeats might be associated with thermodynamically stable secondary structures and yield non-B-form conformations, such as stem-loops or cruciforms. It has been claimed that these short dyad symmetries may play an important role in satDNA repeats as targets for protein binding and thus in satDNA function [12,44,53,66,67,68,69]. Kasinathan and Henikoff [70] have proposed that that cruciform structures formed by dyad symmetries may specify centromeres and that these non-B form DNA configurations in centromeric repeats may facilitate centromere assembly [70,71]. In the two grasshopper species analyzed here, short inverted repeats that might facilitate dyad symmetries and non-B DNA conformations were frequent in both shared and non-shared satDNAs, independently of their organization and chromosomal location. We believe that this property is a simple outcome of stochastic processes of satDNA evolutionary dynamics. Its ubiquity suggests that almost any satDNA can be recruited for functions being dependent on the formation of non-B DNA conformations (see Kasinathan and Henikoff [70]).

SatDNA evolution is a topic of high interest for the scientific community, but the processes and mechanisms have sometimes been confused. Molecular drive was a turnover mechanism suggested by Dover [37,38] as a directional force leading to repeat fixation. It has been the prevalent hypothesis for satDNA evolution due to its apparent explicative power as a mechanism for sequence change, turnover, and concerted evolution. Nonetheless, when applied to satDNA, the presence of arrays on multiple genomic sites makes it impossible, in practice, the fixation of a given repeat. The dependence of CTR on the number and extent of satDNA amplifications in *O. decorus* suggests that molecular drive mainly operates through satDNA amplification and is thus a mutational force (e.g. tandem duplication by means of unequal crossing-over). However, the reach of satDNA amplification is limited to changes in the relative abundances of the pre-existing sequence variants for a given family, most frequently leading to incomplete turnovers. A good way to visualize the role of molecular drive (or amplification) in satDNA evolution is through repeat landscapes for families consisting of several subfamilies showing platykurtic curves (i.e. with low abundance and high divergence) and one or two subfamilies displaying leptokurtic distributions (i.e. with high abundance and low divergence) (see Fig. 2 and Additional file 2: Fig. S1), the latter being those sequences that acquire relevance through satDNA amplification. The comparison of orthologous satDNA pairs between species thus reveal that satDNA amplification implies molecular drive or drift at intra- and inter-specific levels, respectively.

The high or low degree of homogenization for a given satDNA is inversely proportional to the time since the last amplification. It thus depends on i) the neutral mutation rate introducing new sequence variants (increasing intra-specific divergence) and ii) the rate of satDNA amplification, implying partial turnovers that promote sequence variants that become new subfamilies. As satDNA amplification for orthologous satDNA families is independent in relative species, it behaves as an inter-specific drifting mechanism. This dual role of satDNA amplification as the major homogenizing force at the intraspecific level and as the principal driver for interspecific sequence divergence, forced by reproductive barriers, inevitably leads to the concerted evolution pattern. In fact, 16 pairs of orthologous satDNAs met this pattern, with only four showing a non-concerted one. Remarkably, these exceptions coincided with the absence of major amplifications in *O. decorus* satDNAs that remain at low abundance. This kind of variation can persist for long in the absence of (homogenizing) amplification events [72]. Therefore, concerted evolution should be a reasonable consequence of the stochastic nature of satDNA evolution, while exceptional non-concerted patterns can result from differential amplifications among species. Other exceptions can result from satDNA homology with TEs, as was the case for LmiSat02-176, whose homology with Helitron might have biased the calculation of intraspecific divergence. Other explanations have been raised as possible causes for non-concerted evolution patterns, such as the effect of location, organization, and repeat-copy number [55,72,73], population and evolutionary factors [29,33,75,76,77], biological factors [68,77], or functional constraints [32].

We have shown here that concerted evolution is a pattern emerging from satDNA amplification due to the resulting homogenization at intraspecific level and diversification at interspecific level. To visualize this relationship, think about two species recently emerged from a common ancestor. Their satDNA libraries are almost identical at interspecific level but both retain the ancestral polymorphism at intraspecific level. This situation would imply, for each OSF, ILibS values next to 1 and CEI<0 since divergence would be higher at intra-than inter-specific level. As time goes by and mutation and drift operate, ILibS will decrease and CEI will increase as new mutations occur independently in both species. In absence of satDNA amplification, mutation and drift would lead satDNA towards concerted evolution by increasing interspecific divergence, although this process would be slow. However, the pathway to concerted evolution would be paved away by satDNA amplification as the resulting homogenization would reach CEI>0 values (by sharply decreasing intraspecific divergence) when ILibS would decrease below a threshold which, in the case of *O. decorus* and *L. migratoria*, lies between 0.84 and 0.88. The fact that this threshold is so close to 1 reinforces the idea that concerted evolution is an unavoidable property fastly emerging from satDNA amplification. In fact, the four satDNA families which in *O. decorus* showed signs of non-concerted evolution showed low levels of homogenization (RPS between 0.29 and 0.40) and high values of ILibS (0.88-1), presumably due to the low level of amplification of these four satDNAs in this species. Taken together, our results indicate that concerted evolution is a state of interspecific diversification of the satDNA library, reached below a given ILibS threshold, which is fastly promoted by satDNA amplification.

### A model for satDNA evolution

Considering all findings derived from the quantitative analysis of 114 satDNAs in *O. decorus* and *L. migratoria*, we suggest the following model for satDNA evolution (Fig. 6). Intragenomic changes are mainly stochastic, implying that satDNA families mainly evolve under the domain of mutation and drift. SatDNA arises from any tandem duplication yielding at least two monomers. Subsequent unequal crossover is the main source for longer arrays with the consequent increase in tandem structure. This tandem duplication is one of the two classes of mutation operating on satDNA. The other is point mutation increasing divergence among the different monomers composing the whole set of satDNA sequences belonging to a given family. When tandem duplication occurs massively during a short time, it constitutes an **<u>amplification</u>** event that decreases intra-specific divergence (i.e., increases homogenization as measured by RPS) by adding a high number of repeats showing identical sequence. Next, intra-specific divergence will grow across years by the incidence of point mutations, inevitably leading to the **<u>degeneration</u>** of the satDNA sequence unless new amplifications occur. This is characterized by a temporal decrease of RPS and kurtosis and an increase of DIVPEAK as family sequences became more and more divergent. From time to time, some monomers will lose their identity as members of a given satDNA family (reaching identities lower than 80%) or even as members of the same superfamily (with no recognizable homology). This process may shorten long arrays into pieces, thus decreasing TSI and, finally, the satDNA may fade away across time.

**Figure 6.**
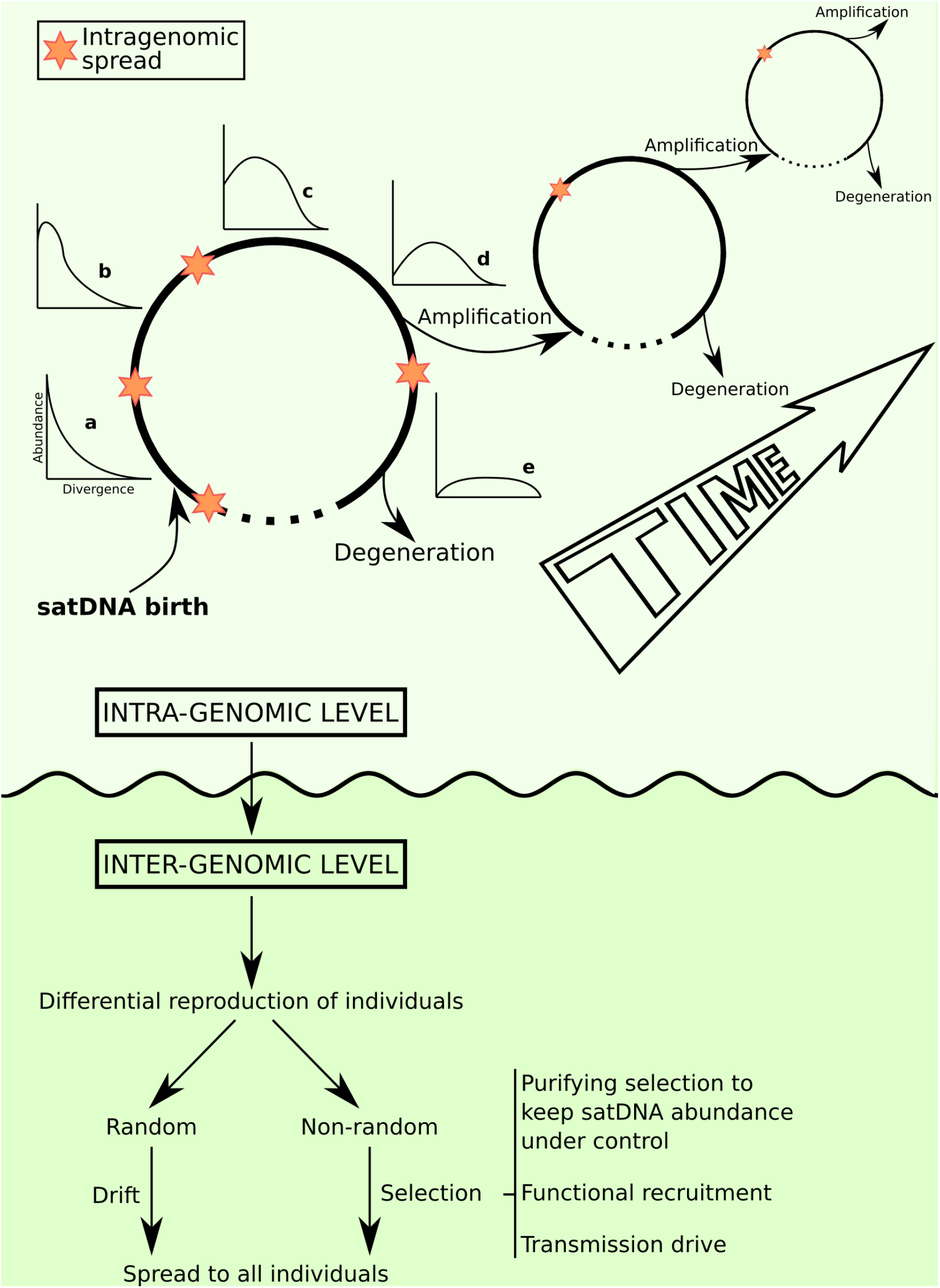
A model of satDNA evolution. We consider that evolutionary events are rather different at intra- and inter-genomic levels. At intra-genomic level, tandem duplication gives birth to a new tandem repeat and its reiteration yields many copies of identical non-coding sequences (satDNA amplification). The newly amplified satDNA displays RLs sharply leptokurtic (a). As time goes by, point mutation increases divergence among the amplified sequences and the curve progressively is flattened (b-e) and DIVPEAK (i.e. the divergence value showing the higher abundance) increases (i.e. the peak moves to the right in the a-e graphs). At any moment of this first amplification-degeneration cycle, another sequence undergoes amplification and begins a new cycle. This sets the satDNA family farther from degeneration and extinction because its average divergence decreases and now predominates a newly amplified subfamily with leptokurtic RL (we represent here three successive cycles of amplification; note that the differences in size among cycles are to facilitate drawing and have nothing to do with amplification level). In parallel, an intra-genomic spread of the satDNA can occur at higher or lower extent (brown stars). A conceivable exit of these cycles is satDNA degeneration, when homology with the original sequence is lost. At inter-genomic level, individual reproduction will mark the destiny of the different satDNA sequences in populations. When reproduction is differential, albeit random (drift) or non-random (selection), some sequences may become prevalent above others. At this respect, the mutational-hazard hypothesis is applicable to explain the limits to purifying selection in some species showing extremely high abundance of satDNA. Finally, we cannot rule out that, in some case, transmission drive could help satDNA to prosper and, even that positive selection may recruit satDNA for important functions, such as telomeric or centromeric functions.

Each new amplification event drives a satDNA family away from degeneration (by promoting that a given subfamily shows the highest abundance and homogenization), after which new point mutations will drive it towards degeneration again, and even complete disappearance if new amplifications do not take place. In summary, we suggest that satDNA undergoes recursive cycles of amplification-degeneration that may keep them in the genome for a long time. During this time, they can integrate into longer repeat units or higher-order structures [79,80], or else disappear through sequence degeneration and/or unequal crossover. The fact that short satDNAs degenerate faster than the longer ones (see above) suggests that their cycle is usually shorter than that of long satDNAs, partly explaining why many short satDNAs show high K2P divergence and platykurtic distribution. For instance, LmiSat10-9 is made of monomers of only 9 bp and is not found in Ode. Even if it would have been present in the common ancestor, it is doubtful that it would have remained for 22.8 Ma in both species without losing identity in at least one of them. In fact, there seems to be a minimum monomer length for homology conservation in these two species, which was 57 bp (LmiSat27-57 and OdeSat41-75). Alternatively, a satDNA formed by repeats of only 9 bp could have arisen *de novo*, by chance, in the gigantic genome of *L. migratoria* [22].

In addition to all former intragenomic events, satDNA frequently undergoes spread among chromosomes. Transposition and replication of extrachromosomal circles of tandem repeats, by the rolling-circle mechanism, followed by reinsertion of replicated arrays, have been postulated as the main mechanisms for the amplification and spread of satDNA families and is supported by indirect [43,81] or direct [14,15] evidence.

At intergenomic (population) level, the only conceivable way to spread an amplification event (occurred in a single individual) is through differential reproduction, as we believe that the molecular drive mechanism suggested by Dover [37,38] as a non-selective fixing force even at the population level, is circumscribed at the intragenomic level. Differential reproduction can occur at random, i.e., by genetic drift, or non-random, i.e., through selection. The latter may be negative, setting up an upper limit to the amount of satDNA tolerable by a genome. Purifying selection, mutation and drift are the drivers in the mutational-hazard (MH) hypothesis [82,83], which suggests that the efficacy of purifying selection is impaired by genetic drift in small populations. This is especially applicable to satDNA, where CTR is highly variable among families (intragenomically). The fact that all satDNA families within a genome have been submitted to the same demographic changes at population level (excepting the differences due to sex linkage) means that purifying selection appears to set few limits to the variation in nucleotide substitution rate among satDNA families. Interestingly, 18 out of 20 shared satDNA families in *L. migratoria* showed amplification events giving rise to FISH bands, whereas only six out of their 14 orthologous families in *O. decorus* did it. This reveals that many of these OSFs have shown highly different evolutionary paths in both species. Based on the MH hypothesis, we may speculate that the extreme demographic changes associated with locust outbreaks in *L. migratoria* might have helped to spread individual satDNA sequences at the population level during the extreme bottlenecks that characterize the solitary phase and subsequent population expansions during the gregarious one. This issue needs further research, including quantitative population analyses of every satDNA family in this species.

In addition, selection can operate positively through non-phenotypic (i.e., meiotic drive) or phenotypic (functional recruitment) effects, as is the case for centromeric and telomeric repeats. The latter is the extreme example of functional recruitment since the repeat is actively homogenized by an RNA-protein complex (telomerase) coded by the genome. Centromeric satDNA in primates resembles this kind of recruitment as another gene (CENPB) is involved in the organization of centromeric satDNA [62,63,64,65].

Our model is an extension of the models devised in the ‘70s and ‘80s [4,5,6,7,8,9,10,11], with some more emphasis on the intragenomic level, and under the light of the MH hypothesis [82,83]. Briefly, amplification is the homogenizing force of satDNA whereas point mutation causes sequence degeneration, with both forces acting recursively. We believe that our model brings about some essential term clarifications. For instance, Escudeiro et al. [84] recently suggested a model of satDNA evolution in bovids consisting of three stages, namely amplification, degeneration (deduced from high satDNA similarity between some species and low between others) and homogenization (high sequence identity among all species). These authors thus claimed for degeneration and homogenization as if they were inter-specific processes. However, in our model, both processes are intragenomic (i.e., intra-specific) resulting from satDNA amplification and point mutation, respectively, whereas inter-specific homogenization or degeneration is highly unlikely under contingent evolution. In fact, homogenization to an identical sequence in several species could only be achieved by functional (selective) recruit, as that occurred for the telomeric DNA repeat.

Finally, the paralogous nature of the satDNA library implies that its diversification between species may show high levels of incomplete library sorting, and this may be a problem for the use of satDNA for phylogenetical purposes beyond satDNA evolution itself. However, the pathway followed by an ancestor satDNA library after speciation can be monitored by satellitome comparison, as shown here for *O. decorus* and *L. migratoria*. A new body of research is taking form recently about contingency and determinism in evolution [46], trying to answer Gould’s question on whether evolutionary trajectories are repeatable [85]. In this respect, satellitome evolution is a natural “parallel replay experiment” able to show many properties of contingent evolution, as the initially identical libraries in the ancestor undergo independent evolution after speciation reaching a high diversity of outcomes among different OSFs. Within species, the environment (at both intragenomic and population levels) is the same for all satDNA families (except for genomic location and organization), but the pathway followed by each of them is highly variable: some families show consensus sequences being highly similar to those in the other species, thus showing high ILibS, whereas others are completely sorted between species, and still others are unrecognizable between species because they have arisen *de novo* in one species or else they have undergone so many sequence changes that have lost homology between species. In analogy with Blount et al. [46] claiming at ecological level, the evolutionary trajectory followed by each OSF in the satellitomes of two separate species is mainly influenced by stochastic processes (i.e. mutation and drift), most likely reaching different outcomes even when both species satellitomes started from the same state in the ancestor and the different OSFs evolved under almost identical conditions at intragenomic level. Therefore, the satellitome is a good example of contingent evolution supporting that “disparate outcomes become more likely as the footprint of history grows deeper” [46]. A rough estimate of the minimal degree of contingent evolution in the *O. decorus* and *L. migratoria* satellitomes can be obtained from the 20 orthologous satDNA pairs used here to estimate CTR. As Table 2 shows, only two of them showed identity higher than 95%: OdeSat17-176/LmiSat02-176 showing a single nucleotide difference in their consensus sequences, and OdeSat59-185/LmiSat01A-185 showing two differences. The first pair showed homology with Helitron TEs which could have biased identity calculations, and the second one appears to have little to do with functional conservation (as explained above). Even assuming that these two cases are adaptive convergences (which is unlikely), we can estimate that satDNA evolution in these species was at least 90% contingent.

The comparison of the satellitomes in two grasshopper species belonging to the subfamily Oedipodinae has allowed us to develop several indices that have proven to be highly useful in the joint analysis of tens of different satDNA families. These were TSI (tandem structure index), RPS (relative peak size) and kurtosis of the repeat landscape distribution as homogenization indices, DIVPEAK as an index of degeneration, CEI as an index of concerted evolution, CTR for consensus turnover rate, and IlibS for incomplete library sorting. However, the main shortcoming of our present analysis was the impossibility to ascertain whether those satDNA families showing no sequence homology between these two species (i.e., non-shared satDNAs) arose *de novo* in one of the species or else they had degenerated in one species but not in the other. To solve this problem, it will be necessary to analyze many species belonging to the same taxonomical group and thus sharing a given satDNA library. We are now sequencing other oedipodine species to perform a multispecies satellitome comparison in the hope that it will allow a better classification of the non-shared satDNA families into *de novo* and partly extinct ones.

## Conclusions

The analysis of the satellitomes of two species of grasshoppers separated by 22.8 Ma of independent evolution has revealed that one-third of the nearly 60 satDNA families found in each species showed sequence similarity to be considered orthologous and thus descended from their last common ancestor. SatDNA turnover at the level of consensus sequences (CTR) showed a range of variation up to two orders of magnitude among orthologous superfamilies. The use of new satDNA parameters allowing to quantify tandem structure (TSI), homogenization (RPS), degeneration (DIVPEAK), concerted evolution (CEI) and incomplete library sorting (ILibS) showed that satDNA amplification has a dual role by increasing homogenization at intra-specific level and diversification at inter-specific level, thus being a molecular driver unavoidably leading to concerted evolution. Most orthologous pairs of satDNAs analyzed in these species showed the concerted pattern of evolution. The causes for the four non-concerted evolution cases were identified as poor amplification in *O. decorus*. The highest levels of concerted evolution were found for satDNAs displaying long repeat units, high levels of homogenization and FISH bands. These results led us to put forward a general model for satDNA evolution, which updates past models with new empirical data and new statistical approaches to quantify key aspects of variation in satDNA dynamics. We also provide a renewed view of the Library Hypothesis by which a satDNA library begins a new divergence process with each cladogenetic event, during which some satDNA families can disappear whereas other can form *de novo*. The contingent nature of satDNA evolution will make unpredictable the precise set of satDNAs present in each species, some of which will be shared with other species and others will not.

## Methods

### Materials and sequencing

We collected 21 males of the grasshopper *Oedaleus decorus* in Cortijo Shambala (Sierra Nevada, Granada, Spain; 36.96111 N, 3.33583 W) on 6 July 2015. They were anaesthetized with ethyl-acetate vapours prior to dissection, and testes were fixed in 3:1 ethanol-acetic acid and stored at 4°C for subsequent fluorescent in situ hybridization (FISH) analysis. Body remains were immersed in liquid nitrogen and stored at −80 °C for molecular analysis and DNA sequencing. We then extracted genomic DNA from a hind leg from one male, using the GenElute Mammalian Genomic DNA Miniprep kit (Sigma). Next we sent the purified DNA to Macrogen Inc. (South Korea) who built a genomic library with ∼180 bp insert size, using the Illumina Truseq nano DNA kit, and sequenced it in an Illumina HiSeq2000 platform (2×101 nt) yielding about 9 Gb of reads. We deposited this library in the Sequence Read Archive (SRA) under accession number SRR9649806 [86].

For the *Locusta migratoria* satellitome, we used the results generated in Ruiz-Ruano et al. [22], including some new analyses of the same Illumina libraries obtained from a Spanish individual lacking B chromosomes (SRA library SRR2911427 [87]), satDNA FISH location, and their consensus sequences (GenBank accession numbers KU056702–KU056808). During these new analyses, we detected a previous mistake in the assembly of the LmiSat01A-193 subfamily, consisting of a false tandem duplication of 8 nt in the consensus monomer. We amended this mistake and renamed the (new) sequence as LmiSat01A-185 (GenBank accession number KU056702.2). We thus performed a new analysis of abundance and divergence for the whole satellitome, considering this modification that implied only slight changes.

In addition, we generated an Oxford Nanopore library for *L. migratoria* using the MinION system with a flow cell version R9. We constructed the library using 5 μg of DNA without fragmentation step applying the the Nanopore Genomic Kit version SQK-LSK108 and the CleanNGS magnetic beads for washes. After applying the localbase-calling program from Nanopore, we got 63,346 reads summing up 130 Mb (∼0.02x of coverage).

### Bioinformatic and sequence analyses

We characterized the *O. decorus* satellitome applying the satMiner protocol [22]. Briefly, this protocol begins with a run of RepeatExplorer [88] and the elimination of homologous reads with Deconseq [89] to perform a new round of RepeatExplorer with the remaining reads. We started with 100,000 read pairs and performed five additional rounds, subsequently duplicating the number of read pairs. Then we identified clusters in each RepeatExplorer round showing spherical or ring-shaped graphs, which are typical for satDNA. We checked the structure of their contigs with a dot-plot using Geneious v4.8.5 [90] to test if they were tandemly repeated, and only those that met this condition were considered as satDNA. Every satDNA family was named with three letters alluding to species name (*L. migratoria* or *O. decorus*) followed by “Sat”, a catalogue number (in decreasing order of abundance) and monomer length, following our previous suggestion in Ruiz-Ruano et al. [22]. For instance, the most abundant satDNA families in the two species analyzed here were LmiSat01-185 and OdeSat01-287. The different subfamilies within a same family were alphabetically named with capital letters in order of decreasing abundance.

Considering their level of sequence identity, we classified every collection of homologous sequences into subfamilies (identity>95%), families (>80%), and superfamilies (>40%). Next, we randomly selected 5 million read pairs with SeqTK (https://github.com/lh3/seqtk) and aligned them against the reference sequences with RepeatMasker v4.0.5 [91]. With these results, we estimated total abundance and average divergence and generated a repeat landscape. Finally, we numbered the satellite families in descending order of abundance. We deposited sequences for satellite DNAs characterized in *O. decorus* in GenBank with accession numbers MT009035-MT009125.

We then searched for homology between *L. migratoria* and *O. decorus* satellitomes with the rm_homolgy script [22] that makes all-to-all alignments with RepeatMasker [91]. We aligned homologous satellites with Muscle v3.6 [92] implemented in Geneious v4.8.5 [90] and reviewed them manually. Then we generated minimum spanning trees (MST) with Arlequin v3.5 [93] (Excoffier and Lischer 2010) and visualized them with HapStar v0.7 [94]. We used the same alignments to estimate the divergence between satDNA families of *L. migratoria* and *O. decorus*. To estimate a consensus turnover rate (CTR) of satDNA sequences, we performed alignments of consensus sequences using ClustalX [95]. Sequence divergence between species was calculated according to the Kimura two-parameter model (K2P; [96]), using MEGA6 [97]. When orthologous satDNA families were composed of several subfamilies, all consensus sequences from each subfamily were aligned and the average of all pairwise distances between the two species was computed. Finally, CTR was calculated using the CTR= K/2T equation, where T= divergence time between species and K= K2P divergence (Kimura 1980). Turnover rates were estimated considering that the *Oedaleus* and *Locusta* genera split 22.81 Ma [45].

To get some insights on array length, we analyzed our MinION library obtained from *L. migratoria* gDNA (see above). For this purpose, we performed an alignment of these reads against the consensus sequences of the *L. migratoria* satellitome using RepeatMasker [91]. However, due to the lack of resolution at subfamily level due to the high level of sequencing errors in these long reads, we only performed this analysis only for the most abundant subfamily in each family, i.e, that noted with the letter “A”. We then analyzed the length of all arrays found for each family to recorded the maximum array length (MAL) for subsequent analysis. For this purpose, we only considered arrays showing length higher than 1.5 repeat units, i.e. at least dimers, and the observed figures for MAL in the 56 satDNA families analyzed in *L. migratoria* ranged between 62 and 20,180 repeat units. In addition, we considered 3 nt as the maximum inter-array distance to collapse two consecutive TR arrays into a same array, in order to partly counteract the splitting effect of short insertions or deletions due to replication slippage. These calculations were implemented in a custom script (https://github.com/mmarpe/satION/blob/master/dis_bed_max.py).

### Analysis of tandem structure

We developed a method to estimate the degree of tandem structure in satDNA using a pipeline that we made publicly available throughout repository (https://github.com/fjruizruano/SatIntExt). This method is based on scoring the number of Illumina read pairs containing repeat units for a given satDNA family in the two reads (onwards named “homogeneous read pairs”) and the number of read pairs containing such a repeat in only one member of the read pair (onwards named “heterogeneous read pairs”). The proportion of homogeneous read pairs indicates the degree at which a satDNA family is tandemly structured (tandem structure index = TSI). This index underestimates the true value by the equivalent to the half of the number of arrays (since each array has two external units). However, as the number of repeat units is much higher than the number of arrays, we consider that this underestimation may be low at the genomic level. To validate TSI, we analyzed Oxford Nanopore MinION long reads in *L. migratoria*, by annotating all satDNA variants found in them and scoring the number of repeat units constituting the longest array found for each satDNA family. Despite low coverage of the MinION reads, these longest arrays showed significant positive correlation with TSI (Spearman rank correlation: *r_S_*= 0.42, N= 55, t= 3.36, P= 0.001), indicating that TSI is a valid estimator for the degree of tandem structure of satDNA. In addition, we tried to annotate the external read of every heterogeneous read pair with the database of repetitive elements of *L. migratoria* generated in Ruiz-Ruano et al. [98] with RepeatMasker. Thus, we found homology of the elements adjacent to the satDNA arrays with satDNAs, transposable elements, rDNAs, snDNAs, tRNAs, histones, mitochondrial DNA and unknown elements in some read pairs, and counted the number of occurrences. This analysis is also integrated in the above-mentioned pipeline.

### Homogenization and degeneration indices

SatDNA homogenization, i.e., the degree of intraspecific similarity between its tandemly structured monomers, is conceptually inverse to average sequence divergence. Therefore, a homogenization index should be negatively correlated with the K2P divergence. Trying to get such an index, we built repeat landscapes for each satDNA subfamily (90 in *O. decorus* and 103 in *L. migratoria*) and searched for divergence peaks, i.e., those divergence values showing the highest abundance in the repeat landscape (DIVPEAK) (Fig. 1). Then, we summed up the abundances of all satDNA sequences at ±2% divergence from the DIVPEAK class to calculate abundance in the 5% peak or PEAK-SIZE (Fig. 1). The logic was to get a collection of sequences diverging 5% or less to the consensus sequence, thus coinciding with our criterion to define subfamilies, as they probably derived from the same amplification event (see Ruiz-Ruano et al. [22] for details). Finally, we calculated relative peak size (RPS) as the quotient between PEAK-SIZE and total abundance (see Fig. 1), which measures the proportion of repeat units being part of the last amplification event. To calculate RPS at the family level in those families showing two or more subfamilies, we followed the same procedure including all subfamily satDNA sequences, so that each subfamily weighted in proportion to its abundance. RPS serves as an index of homogenization because it is expected to increase with satDNA amplification, as the new units derived from tandem duplication will initially show identical sequences, thus increasing global identity. DIVPEAK serves as an index of degeneration because it will increase by mutation accumulation and is thus proportional to the time passed since the last amplification. Specifically, DIVPEAK is the value of divergence (from 0% onwards) at which a given satDNA shows its maximum abundance, and increases when mutational decay move its abundance peak away from complete homogenization (divergence=0) where it arrived after its last major amplification event. The values for average divergence, total abundance, maximum abundance, maximum divergence, RPS and DIVPEAK for every satDNA family were estimated from with a custom script using the divsum files from RepeatMasker (https://github.com/fjruizruano/SatIntExt/blob/main/divsum_stats.py).

### Concerted evolution index and incomplete library sorting

We calculated the divergence at intra- (K2P_intra_) and inter-specific (K2P_inter_) levels for the 20 pairs of orthologous satDNA families, and calculated an index of concerted evolution (CEI) as log2 the K2P_inter_/K2P_intra_ quotient.

The comparative analysis of RLs and MSTs revealed that the observed differences between OSFs in CTR were due to the state of library sorting between species. On this basis, we observed that the OSF showing the highest CTR was that showing a best separation between species for all families and subfamilies of satDNA. We then gave 1 to the sorting state of this OSF and then divided all CTR values by this maxCTR to obtain an index of the relative sorting for each OSF. One minus the obtained value thus indicated the degree of incomplete library sorting (ILibS) for each OSF.

### Analysis of conserved motifs and curvature

We analyzed the consensus sequences of shared and non-shared satDNAs between the two species looking for functional signatures. We used the ETANDEM, EINVERTED, and PALINDROME programs from the EMBOSS suite of bioinformatics tools [99] for the detection of internal repeats (direct or inverted) and palindromes. Short internal direct repeats indicate the presence of functional motifs within the satDNA repeats. Dyad symmetries, many of them associated with thermodynamically stable secondary structures, are predicted to adopt non-B DNA conformations, such as stem-loops or cruciforms, which might have a role as targets for protein binding. Thus, as an additional test on the propensity to form non-B DNA conformations, we checked all satDNA families using the Mfold web server (http://www.unafold.org/mfold/applications/rna-folding-form-v2.php) for nucleic acid folding prediction [100], estimating Gibbs free energy (dG) of the predicted secondary structures [101]. We also checked the consensus sequences of both types of satDNAs for sequence-dependent bendability/curvature propensity of repeats. We produced the bendability/curvature propensity plots with the bend.it server at http://pongor.itk.ppke.hu/dna/bend_it.html#/bendit_intro [102], using the DNase I based bendability parameters of Brukner et al. [103] and the consensus bendability scale [104]. Finally, we used the sliding windows option of the DnaSP v.5.10 program [105] for the analysis of nucleotide diversity (π) per position for every shared satDNA in order to detect DNA conserved motifs. For this, we use multiple alignments of several dozens of monomer repeats selected per each satDNA.

### Chromosomal location of the *O. decorus* satDNAs

To compare the chromosomal location of orthologous satDNA families in these species, we performed fluorescent in situ hybridization (FISH) for 14 satDNA families in *O. decorus* which showed sequence homology with 20 families in *L. migratoria*. For this purpose, we designed divergent primers for these 14 satDNA families in *O. decorus* using Primer3 [106] with a Tm ∼60 °C, to generate FISH probes as described in Cabrero et al. [107] and Ruiz-Ruano et al. [22].

### Statistical analysis

To investigate distribution fitting of RPS and DIVPEAK, we used the chi-square test, and the normality of other variable distributions was tested by the Shapiro-Wilks test, and, when this condition was not met, we used the non-parametric Spearman rank correlation test. In the case of turnover rate, we performed forward stepwise multiple regression to analyze its dependence on other variables. In this case, we calculated variance inflation factors (VIFs) to test for multicolinearity, and the fit of standardized residuals of this regression to a normal distribution was tested by means of the Shapiro-Wilks test. All these analyses were performed using the Statistica software (Statsoft Inc.). Two-group comparisons were performed by the Gardner-Altman estimation plot method devised by Ho et al. [108] following the design in Gardner and Altman [109], as implemented in https://www.estimationstats.com. This analysis calculates the effect size by the mean difference between groups, for independent samples, or else by the paired mean difference in case of paired samples. The effect size is then evaluated by the 95% confidence interval (95% CI) and whether it includes or not the zero value. Contingency tests were performed by the RXC program, which employs the Metropolis algorithm to obtain an unbiased estimate of the exact p-value [110]. In all cases 20 batches of 2,500 replicates were performed.

## Abbreviations

B-pattern: Banded pattern (pattern in FISH analyses)
CEI: Concerted Evolution Index
CI: Confidence Interval
CTR: Consensus Turnover Rate
dG: Gibbs free energy
DIVPEAK: Divergence Peak
FISH: Fluorescence *In Situ* Hybridization
ILibS: Incomplete Library Sorting
K2P: Kimura Two-Parameter (substitution model)
Lmi: *Locusta migratoria*
NS-pattern: No signal pattern (in FISH analyses)
MAL: Maximum Array Length (observed in MinIon reads of *L. migratoria*)
MST: Minimum Spanning Tree
Ode: *Oedaleus decorus*
OSF: Orthologous Superfamily
RL: Repeat Landscape
RPS: Relative peak size
RUL: Repeat Unit Length
satDNA: satellite DNA
SF: Superfamily
TSI: Tandem Structure Index
VIF: Variance inflation factors

## Declarations

### Ethics approval and consent to participate

Not applicable.

### Consent for publication

Not applicable.

### Availability of data and materials

The Illumina libraries used for this article are available in the Sequence Read Archive (SRA) with accession numbers SRR9649806 [86] and SRR2911427 [87]. Main data generated or analyzed during this study are included in this published article and its supplementary information files.

### Competing interests

The authors declare no competing interests.

### Funding

FJRR was also supported by a postdoctoral fellowship from Sven och Lilly Lawskis fond (Sweden) and a Marie Skłodowska-Curie Individual Fellowship (grant agreement 875732, European Union).

## Acknowledgments

Not applicable.

## Authors’ contributions

Conceptualization: JPMC, JC, MDLL, MMP, FP, MAGR, FJRR; experimental design: JPMC, JC, MDLL, MMP, FP, MAGR, FJRR; sampling: JPMC and JC; cytogenetic analyses: JPMC, JC, MDLL; data analysis: JPMC, MMP, MAGR, FJRR. All authors read and approved the manuscript.

## Supplementary Information

### *Additional file 1 (.xls format): Tables S1-S4

**Table S1.** Molecular and cytological properties of the satellitomes in *Oedaleus decorus* (Ode) and *Locusta migratoria* (Lmig). Note that telomeric DNA was also numbered in both species (no. 13 and 7, respectively) but are omited here because they were not considered for this paper analyses. RUL= Repeat unit length. TSI= Tandem structure index. SF= Superfamily. RPS= Relative peak size. DIVPEAK= Divergence peak. MAL= Maximum array length observed in MinIon reads of *L. migratoria*. FISH= FISH pattern (B= banded, NS= No signal). Local= Localization (p= proximal, i= interstitial, d= distal). Motifs= Conserved motifs in the DNA sequence (0= Yes, 1= No). Curvature= Propensity to adquire stable structures (0= Yes, 1= No). dG= Gibbs gree energy of the predicted secondary structure.

**Table S2.** Homology between satDNA families found in *O. decorus* and *L. migratoria*. OSF= Orthologous superfamily. Those families chosen for comparisons between orthologous pairs are noted in bold-type letter.

**Table S3.** Total number of external reads for each satellite family in *O. decorus* (Ode) and *L. migratoria* (Lmig) and its annotation. TSI= Tandem Structure Index.

**Table S4.** Characteristics of the orthologous satDNA families analyzed in *O. decorus* (14) and *L. migratoria* (20). Each row includes one Ode and one Lmi satDNA families showing homology. Note that some Ode families showed homology with two or three Lmi ones. OSF= Orthologous superfamily, sf= number of subfamilies, SF= superfamily name, FISH= FISH pattern (B= banded, NS= no signal), RUL=Repeat unit length (bp), A+T= % A+T content, abun= abundance (% of the genome), div= divergence (%), peak_size= abundance of the 5% divergence classes around DIVPEAK, RPS= Relative peak size, DP= DIVPEAK, kur= kurtosis of repeat landscape distribution, TSI= Tandem structure index, dG= Free energy of repeat unit sequence, MAL= Maximum array length observed in MinIon reads of *L. migratoria*, CEI= Concerted evolution index (L= *L. migratoria*, O= *O. decorus*), Intid= Interspecific sequence identity (%), Intdiv= Interspecific divergence, CTR= Consensus turnover rate, ILibS= Incomplete library sorting. Negative CEI values and Int_id>95% are remarked in bold type letter.

### *Additional file 2 (.tif format): Figure S1

**Figure S1.** Repeat landscape (RL) and minimum spanning tree (MST) of three orthologous superfamilies of satellite DNA in O. decorus and L. migratoria (OSF04, OSF05 and OSF07). a) RLs showed that OSF04 showed large peaks of amplification in both species but CTR values ranged between 1.16 and 1.6, presumably due to the incomplete library sorting (ILibS) evidenced by the MST (note how OdeSat32A and LmiSat51A connect with both species’ sequences). b) OSF05 showed high CTR values, large amplification peaks in both species and ILibS for only OdeSat22C, which was the only sequence connected with sequences from both species. c) OSF07 showed the lowest CTR values and showed very small amplification peaks for OdeSat58 (green curves in the RL on the left) and higher ILibS, with three sequences being connected with both species’ sequences (LmiSat45-274, LmiSat28A-263 and OdeSat58A-265).

### * Additional file 3 (.xls format): Dataset S1

**Dataset 1a.** Data from the *Oedaleus decorus* repeat landscape indicating genomic abundance for each satellite DNA family and divergence interval.

**Dataset 1b.** Data from the *Locusta migratoria* repeat landscape indicating genomic abundance for each satellite DNA family and divergence interval.

## Notes

### Competing Interest Statement

The authors have declared no competing interest.

