## Supplementary figures and images for "Satellitome comparison of two oedipodine grasshoppers highlights the contingent nature of satellite DNA evolution"

### Additional file 2, Supplementary Fig. S1

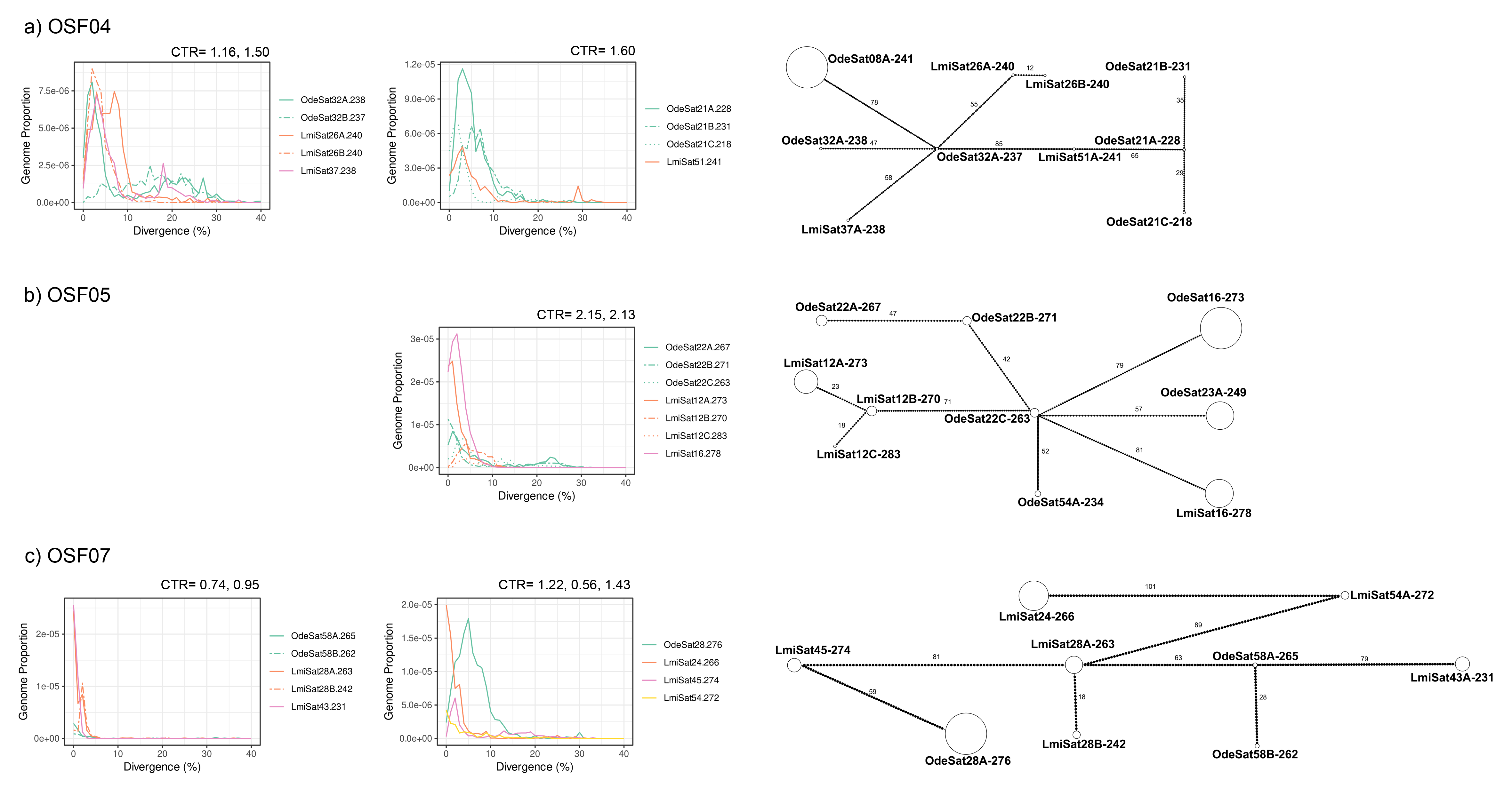
